# Transcription and splicing dynamics during early *Drosophila* development

**DOI:** 10.1101/2020.11.05.367888

**Authors:** Pedro Prudêncio, Rosina Savisaar, Kenny Rebelo, Rui Gonçalo Martinho, Maria Carmo-Fonseca

## Abstract

Widespread co-transcriptional splicing has been demonstrated from yeast to human. However, most studies to date addressing the kinetics of splicing relative to transcription used either *Saccharomyces cerevisiae* or metazoan cultured cell lines. Here, we adapted native elongating transcript sequencing technology (NET-seq) to measure co-transcriptional splicing dynamics during the early developmental stages of *Drosophila melanogaster* embryos. Our results reveal the position of RNA polymerase II (Pol II) when both canonical and recursive splicing occur. We found heterogeneity in splicing dynamics, with some RNAs spliced immediately after intron transcription, whereas for other transcripts no splicing was observed over the first 100 nucleotides of the downstream exon. Introns that show splicing completion before Pol II has reached the end of the downstream exon are necessarily intron-defined. We studied the splicing dynamics of both nascent pre-mRNAs transcribed in the early embryo, which have few and short introns, as well as pre-mRNAs transcribed later in embryonic development, which contain multiple long introns. As expected, we found a relationship between the proportion of spliced reads and intron size. However, intron definition was observed at all intron sizes. We further observed that genes transcribed in the early embryo tend to be isolated in the genome whereas genes transcribed later are often overlapped by a neighboring convergent gene. In isolated genes, transcription termination occurred soon after the polyadenylation site, while in overlapped genes Pol II persisted associated with the DNA template after cleavage and polyadenylation of the nascent transcript. Taken together, our data unravels novel dynamic features of Pol II transcription and splicing in the developing *Drosophila* embryo.

## INTRODUCTION

It is by now largely accepted that splicing can happen, and often does happen, whilst transcription is still in progress (Ameur et al. 2011; Khodor et al. 2011; Beyer and Osheim 1988; Carrillo Oesterreich et al. 2010; Windhager et al. 2012; Nojima et al. 2015; Alpert et al. 2017; Brugiolo† et al. 2013). It is also known that the processes of splicing and transcription are tightly interlinked. The RNA Polymerase II (Pol II) elongation rate can affect exon inclusion (Fong et al. 2014; Aslanzadeh et al. 2018; de la Mata et al. 2003; Maslon et al. 2019), and many of the proteins involved in splicing associate with the Pol II large subunit C-terminal domain (CTD) (Hsin and Manley 2012; Emili et al. 2002; David et al. 2011; Görnemann et al. 2011; Morris and Greenleaf 2000; Nojima et al. 2018). Conversely, splicing may also affect transcription, with evidence suggesting that Pol II slows down at exons (Alexander et al. 2010; Carrillo Oesterreich et al. 2010; Mayer et al. 2015; Veloso et al. 2014; Jonkers et al. 2014), potentially to allow for splicing to complete.

Several studies have addressed the timing of intron excision relative to Pol II elongation. A nascent RNA sequencing study (Carrillo Oesterreich et al. 2016) showed that in *Saccharomyces cerevisiae*, splicing could occur as soon as the 3’ splice site had emerged from the transcription machinery, suggesting that splicing may be completed immediately after intron transcription. Previous attempts to determine the duration of splicing *in vivo* had returned estimates ranging from a few seconds to several minutes (Alpert et al. 2017). The lowest of these estimates, such as the few-second estimate reported in Martin et al. (2013), are consistent with splicing being completed right after the 3’ splice site is transcribed. The most recent breakthroughs in the field followed the development of long-read nascent RNA sequencing technologies (Drexler et al. 2020; Reimer et al. 2021; Sousa-Luís et al. 2021). Collectively, these studies performed in human and *Drosophila* cultured cell lines show that although many introns are immediately excised as soon as the downstream exon emerges from Pol II, a subset remains unspliced and progressively undergoes delayed splicing while Pol II transcribes further (Drexler et al. 2020; Reimer et al. 2021; Sousa-Luís et al. 2021).

Here, we studied the dynamic properties of transcription and co-transcriptional splicing during early stages of development in *Drosophila melanogaster* embryos using Native Elongating Transcript Sequencing (NET-seq) (Churchman and Weissman 2011; Nojima et al. 2015). An advantage of using *Drosophila* embryos is that they are more physiological than cultured cell lines. Moreover, compared to human, the *Drosophila* genome is more compact (Graveley et al. 2011) and thus the coverage of NET-seq reads on intragenic regions is higher. Another advantage of the *Drosophila* model is that genes transcribed initially in the embryo have few and short introns (like yeast genes), whereas genes transcribed later contain multiple long introns (more similar to human genes). An additional feature of *Drosophila* is the presence of many genes with exceptionally long introns that are subdivided by a non-canonical mechanism termed recursive splicing (Duff et al. 2015; Joseph et al. 2018; Pai et al. 2018). For all these reasons, *Drosophila* is an attractive model to study co-transcriptional splicing dynamics.

*Drosophila* early development is characterized by rapid mitotic cycles that lack cytokinesis, resulting in nuclear proliferation in a syncytial cytoplasm (Campos-Ortega 1985; Laver et al. 2015). During these initial mitotic divisions, which impose significant constraints on transcription and splicing (Shermoen and O’Farrell 1991; Rothe et al. 1992; Martinho et al. 2015; Sandler et al. 2018; Kwasnieski et al. 2019; Guilgur et al. 2014), the embryo largely relies on maternally deposited mRNAs. Subsequently, the duration of the cell cycle is progressively expanded, membranes form between the nuclei and segregate them into cells, and the zygotic transcriptome starts to be fully expressed (Campos-Ortega 1985; Yuan et al. 2016; Blythe and Wieschaus 2015). By performing *d*NET-seq in the developing *Drosophila* embryo, we have unravelled novel dynamic features of Pol II transcription and splicing.

## RESULTS

### Native elongating transcript sequencing in *Drosophila* embryos (*d*NET-seq)

Our first task was to adapt the NET-seq technology for use in *Drosophila* embryos (henceforth referred to as *d*NET-seq). The NET-seq technique involves isolation of transcription complexes formed by Pol II, the DNA template and the nascent RNA by immunoprecipitation, without crosslinking (Churchman and Weissman 2011; Nojima et al. 2015). After solubilisation of Pol II complexes under native conditions by extensive micrococcal nuclease (MNase) digestion of isolated native chromatin, elongation complexes were immunoprecipitated using antibodies that specifically recognize different phosphorylation states of the Pol II CTD (Nojima et al. 2015).

We collected embryos at 2-3 hours after fertilization (referred to as early embryos) and 4-6 hours after fertilization (referred to as late embryos) (**Fig 1A**). Analysis of embryos stained with a fluorescent dye to visualize DNA (**Fig 1B**) revealed that early embryos were predominantly in mitotic cycle 14 (stage 5), which is when massive activation of zygotic transcription occurs (Campos-Ortega 1985; Laver et al. 2015). The majority of late embryos were in the late stage of germ-band extension (stage 10; **Fig 1C**), when the embryo trunk (also known as the germ-band) elongates in the antero-posterior axis and narrows in the dorso-ventral axis (Campos-Ortega 1985; Laver et al. 2015).

**Figure 1.**
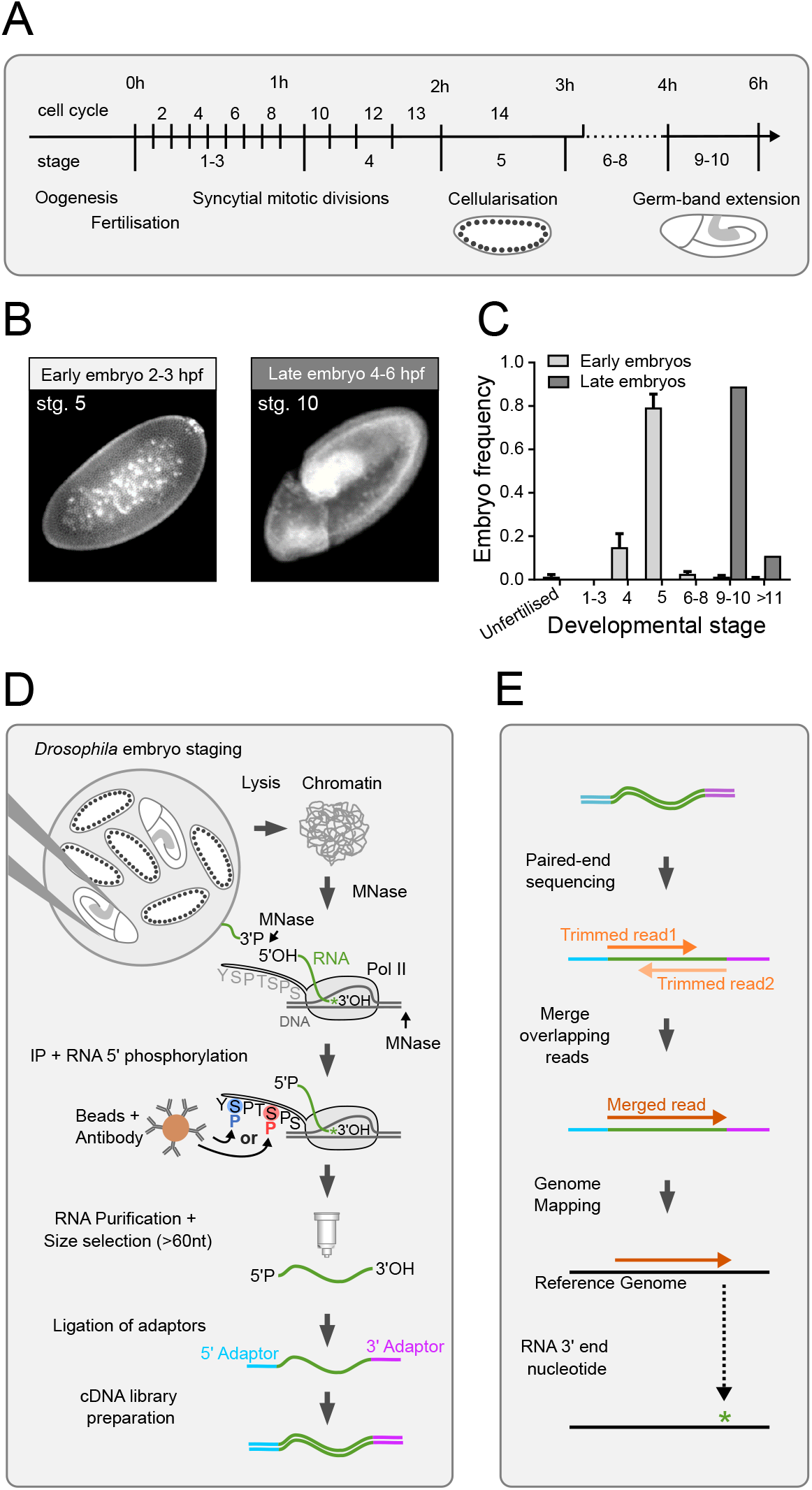
Native elongating transcript sequencing in *Drosophila* embryos. **(A)** Timeline of *Drosophila* early embryonic development, which starts with 13 rapid syncytial mitotic cycles. During interphase of cycle 14, membranes form between the nuclei located at the periphery of the embryo (cellularisation). The new cells start morphogenetic movements leading to elongation of the embryo trunk (germ-band extension). **(B)** Representative images (stained for DNA) of embryos in mitotic cycle 14 (stage 5) and late germ-band expansion (stage 10). **(C)** The graph depicts the developmental stage of embryos sorted into the “early” and “late” groups. Approximately 30000 early embryos and 15000 late embryos were analysed. **(D)** Outline of the *d*NET-seq experimental protocol. **(E)** Outline of *d*NET-seq data analysis.

In adapting mNET-seq to *Drosophila* embryos, we optimized buffers and washing conditions to purify the chromatin fraction from manually sorted embryos, solubilize the transcription complexes with MNase digestion and immunoprecipitate Pol II with antibodies. We used rabbit polyclonal antibodies raised against synthetic peptides of the YSPTSPS repeat of the CTD of the largest Pol II subunit in *Saccharomyces cerevisiae*, phosphorylated at either S5 (ab5131 Abcam) or S2 (ab5095 Abcam). Both antibodies have been extensively used for chromatin immunoprecipitation experiments in *Drosophila melanogaster* (Boija et al. 2017; Vizcaya-Molina et al. 2018; Akhtar et al. 2019; Dahlberg et al. 2015; Lee et al. 2015; Yan et al. 2014; Arzate-Mejía et al. 2020).

To enable directional sequencing, the 5′ hydroxyl (OH) generated by MNase digestion of RNA was first converted to a 5′ phosphate by T4 polynucleotide kinase (**Fig 1D**). RNA was then purified from the immunoprecipitated Pol II complexes and size-selected using an RNA purification kit procedure that combines a unique buffer system with a column technology (see the methods section for more detail). RNAs with a size above 60 nucleotides (nt) were used for subsequent ligation of specific adapters to the 5′ P and 3’ OH ends of each RNA fragment followed by PCR-based preparation of a cDNA library for high-throughput Illumina sequencing (**Fig 1D**). After sequencing, adapter sequences were trimmed and paired-end reads with sequence overlaps were merged into a single read that spans the full length of the original RNA fragment (dark orange; **Fig 1E**). The resulting single reads were aligned to the *Drosophila* reference genome. The nucleotide at the 3’ end of each RNA fragment was identified and its genomic position recorded (asterisk; **Fig 1E**).

Two to three NET-seq libraries were independently prepared from early and late embryos using S5P antibody; three additional libraries were prepared from late embryos using S2P antibody. Each library was sequenced to a high coverage with a read length of 150 bp (Supplemental **Fig S1**; see the methods section for more detail). Experimental reproducibility was demonstrated by strong agreement of uniquely aligned read density between biological replicates prepared with antibodies raised against the CTD phosphorylated on either serine 5 (S5P) or serine 2 (S2P) positions (Supplemental **Fig S1 B-E**). This suggests that both S5P and S2P antibodies recognize the CTD of elongating Pol II in *Drosophila* embryos.

### *d*NET-seq captures splicing intermediates and spliceosomal snRNAs

NET-seq captures not only the final (3’ OH end) nucleotide of nascent RNA but also the 3’ OH end of RNAs that associate with the Pol II elongation complex (Nojima et al. 2015, 2018; Schlackow et al. 2017). Notably, in humans, mNET-seq of Pol II phosphorylated on CTD serine 5 detected splicing intermediates formed by cleavage at the 5′ splice site after the first splicing reaction. The presence of such intermediates manifests as an enrichment of reads whose 3’ ends map precisely to the last nucleotide of an exon. In both early and late *Drosophila* embryos, we indeed observed large peaks of *d*NET-seq/S5P reads mapping to the last nucleotide of spliced exons, as shown for the *kuk* gene (green asterisk; **Fig 2A**). We also detected *d*NET-seq/S5P peaks at the last nucleotide of introns, as shown for the *eEF1alpha1* gene (pink asterisk; **Fig 2B**); enrichment for these reads results from co-immunoprecipitation of released intron lariats after completion of the splicing reaction (**Fig 2B**). In addition, we observed reads corresponding to mature snRNAs engaged in co-transcriptional spliceosome assembly, suggesting that *d*NET-seq was capturing the free 3’ OH ends of the snRNAs. (blue asterisk; **Fig 2C**). We found prominent peaks at the end of spliceosomal U1, U2, U4 and U5 snRNAs. As expected, no peak was detected mapping to the end of the U3 snRNA, which is involved in the processing of pre-rRNA synthesized by Pol I. Noteworthy, we observed an accumulation of *d*NET-seq signal at the end of U6 snRNA (Supplemental **Fig S2A**), contrasting with a lack of peak observed in mammalian cells (Nojima et al. 2018). This is consistent with the finding that most mammalian U6 snRNAs contain a 2’,3’-cyclic phosphate terminal group at the 3’ end, whereas U6 3’ ends in *Drosophila* cells consist of either a cyclic 2’,3’-phosphate, a 3’-phosphate or a 2’,3’-hydroxyl group (Lund and Dahlberg 1992).

**Figure 2.**
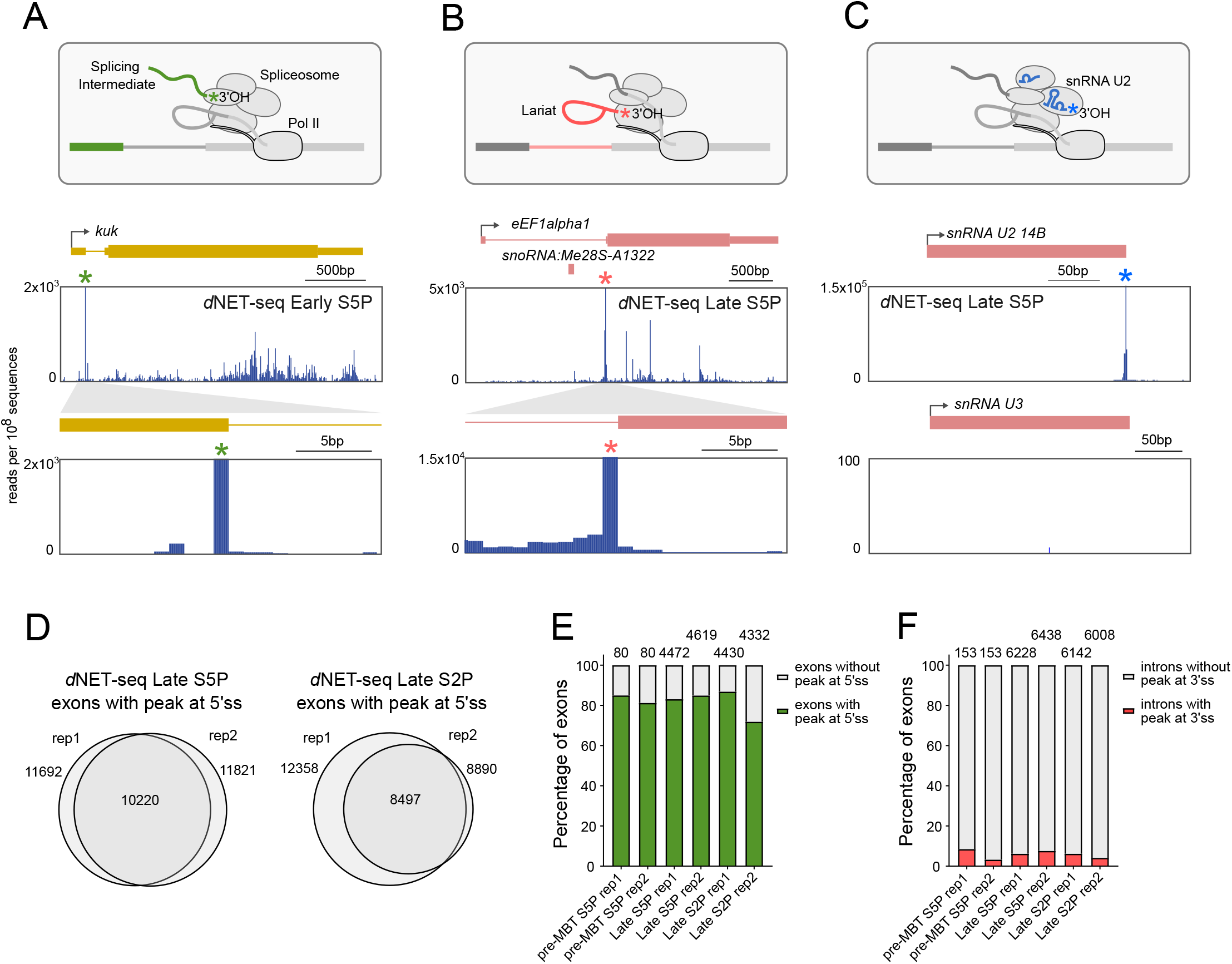
*d*NET-seq captures splicing intermediates and spliceosomal snRNAs. **(A-C)** The diagrams outline the 3’ OH ends generated by co-transcriptional cleavage at the 5’ splice site (**A**), the 3’ splice site (**B**) and the free 3’ OH end of spliceosomal snRNAs (**C**). Below each diagram, *d*NET-seq/S5P profiles over the indicated genes are depicted (data from late embryos). The green asterisk denotes the peak at the end of the exon (**A**). The pink asterisk denotes the peak at the end of the intron (**B**). The blue asterisk denotes the peak at the end of the U2 snRNA gene (**C**). Arrows indicate the direction of transcription. Exons are represented by boxes. Thinner boxes represent UTRs. Introns are represented by lines connecting the exons. (**D**) Comparison (Venn diagrams) of exons with a splicing intermediate peak detected in biological replicates of *d*NET-seq/S5P and *d*NET-seq/S2P libraries. (**E, F**) Frequency of peaks corresponding to splicing intermediates (**E**) and released intron lariats (**F**) in pre-MBT genes and genes expressed in late embryos. Only genes with the highest read density (fourth quartile) were considered.

To quantify how many constitutively spliced exons have *d*NET-seq peaks at the end, we applied an algorithm that finds nucleotides where the NET-seq read density is at least three standard deviations above the transcript mean in a local region defined by 100 bp upstream and downstream (Churchman and Weissman 2011; Prudêncio et al. 2020). Upon analysing replicates of *d*NET-seq/S5P and *d*NET-seq/S2P libraries, we identified over 10,000 exons showing splicing intermediate peaks (**Fig 2D**). As peaks were more frequently detected on exons of genes with higher read density (Supplemental **Fig S2B**), we classified the exons into four groups (quartiles) based on the *d*NET-seq read density of the corresponding gene and restricted the analysis to exons in the fourth quartile, i.e., from genes with the highest read density. The results show that splicing intermediate peaks are detected in approximately 80% of all constitutively spliced exons. We further analysed the so-called pre-MBT (mid-blastula transition) genes, which are the first zygotic genes to become transcriptionally active in early embryos (Chen et al. 2013), and genes expressed in late embryos. We found similar proportions of splicing intermediate peaks associated with pre-MBT and late genes in the S5P and the S2P datasets (**Fig 2E**). We then used the same methodology and the same set of genes to detect peaks at the last intronic nucleotide, corresponding to released intron lariats. Such peaks were detected in less than 10% of introns (**Fig 2F**).

Taken together, these results demonstrate that *d*NET-seq with antibodies that recognize the Pol II CTD phosphorylated at either S5 or S2 positions is capable of detecting splicing intermediates and spliceosomal snRNAs in *Drosophila* embryos, as previously reported in mammalian cells (Nojima et al. 2015; Schlackow et al. 2017; Nojima et al. 2018).

### *d*NET-seq specifically captures nascent RNA

To validate that we were detecting nascent transcription in *Drosophila* embryos, we analysed maternal mRNAs that are transcribed during oogenesis and loaded into the egg (**Fig 3A**). As expected, maternal transcripts such as *bicoid* (**Fig 3B**), *nanos* (Supplemental **Fig S3A**), *gurken* (Supplemental **Fig S3B**) and *Rab32* (Supplemental **Fig S3C**) were detected by RNA-seq in embryos collected 2-3 hours after fertilization, but *d*NET-seq signal over these genes was negligible. However, robust *d*NET-seq/S5P signal was found at the *pumilio* gene (**Fig 3C**). Expression of this gene was considered to be exclusively maternal based on RNA-seq (Lott et al. 2011) and Pol II ChIP-seq data (Chen et al. 2013), but a GRO-seq study detected *pumilio* nascent transcripts in embryos collected 2-2.5 hours post-fertilisation (Saunders et al. 2013). The detection of *pumilio* RNA in early embryos by GRO-seq and *d*NET-seq/S5P highlights the sensitivity of these two techniques in capturing low-level nascent transcripts.

**Figure 3.**
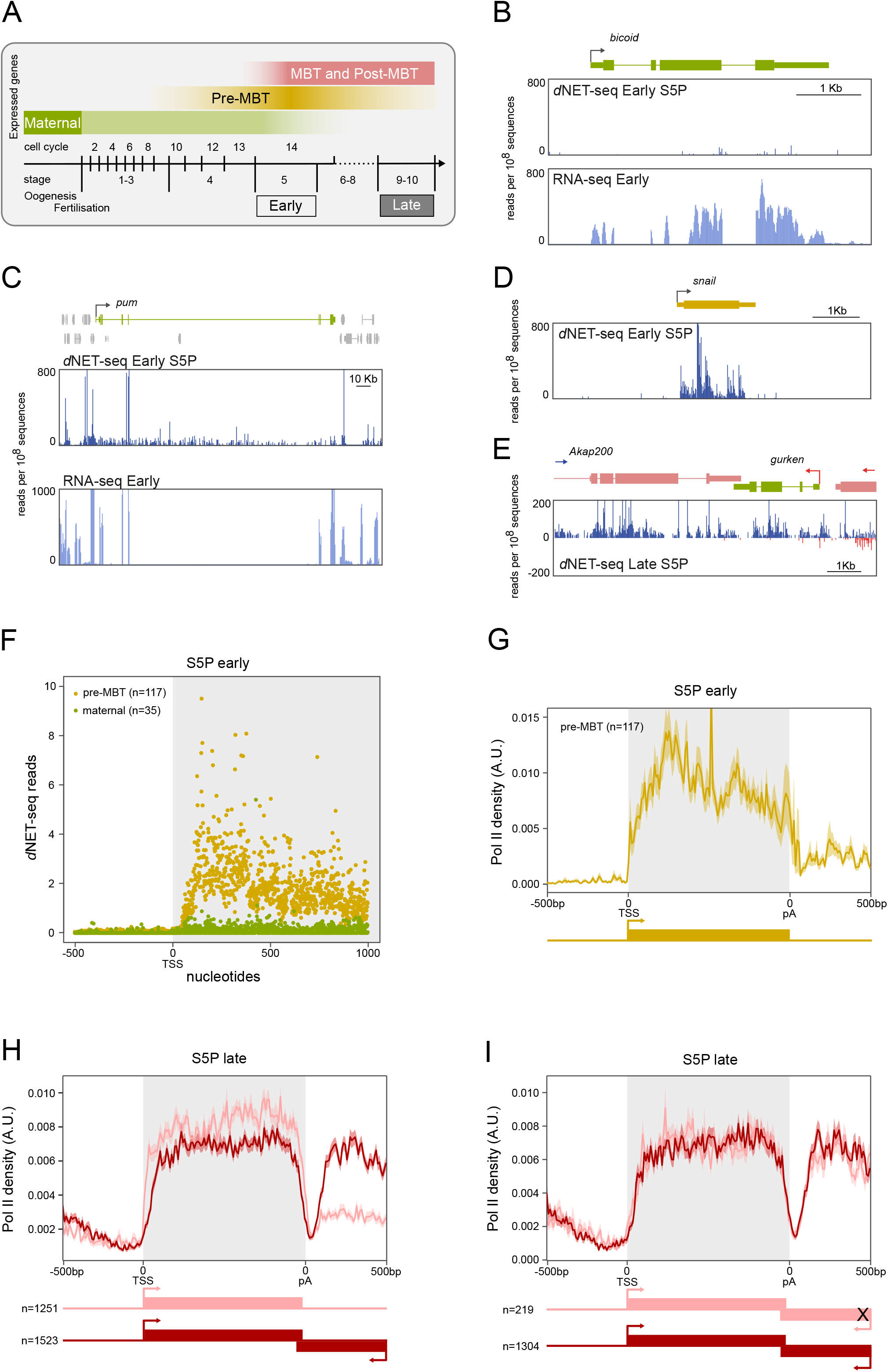
*d*NET-seq profiles in early and late embryos. **(A)** The diagram illustrates the temporal expression of maternal, pre-MBT, MBT, and post-MBT genes during *Drosophila* embryonic development. *d*NET-seq/S5P and RNA-seq profiles over the maternal genes *bicoid* (**B**) and *pumilio (pum)* (**C**), the pre-MBT gene *snail* (**D**), and the post-MBT gene *Akap200* (**E**). Reads that aligned to the positive strand are in blue, and reads that aligned to the negative strand are in red. (**F**) Meta-analysis of mean *d*NET-seq/S5P read density around the transcription start site (TSS) in maternal and pre-MBT genes (replicate 1). **(G-I)** Normalized metagene analysis in arbitrary units (A.U.). The *d*NET-seq/S5P signal is depicted along the normalized gene length (grey background), as well as 500 bp upstream of the transcription start site (TSS) and 500 bp downstream of the polyadenylation (pA) site. **(G)** Pre-MBT genes in early embryos. **(H)** Transcriptionally active genes in late embryos; the signal over genes that have the 3′ UTR overlapped by an antisense gene is depicted in dark red, while the signal over genes with no other genes within 500 bp is depicted in light red. **(I)** Transcriptionally active genes in late embryos; the signal over genes that have the 3′ UTR overlapped by a transcriptionally active antisense gene is depicted in dark red, while the signal over genes with the 3′ UTR overlapped by a transcriptionally inactive antisense gene is depicted in light red.

The stage at which the embryo switches from relying on maternally deposited mRNAs and proteins to undergoing its own transcription is termed the mid-blastula transition (MBT). However, a few genes (known as pre-MBT genes) become transcriptionally active before MBT (Kwasnieski et al. 2019). Strong *d*NET-seq signal was detected in early embryos over the bodies of pre-MBT genes such as *snail* (**Fig 3D**), *fushi tarazu* (**Supplemental Fig S3D**) and *odd skipped* (**Supplemental Fig S3E**). Altogether, we examined 117 previously identified pre-MBT genes and 35 maternal mRNAs (**Fig 3F and Supplemental Fig S3F**). The finding that robust *d*NET-seq/S5P signal was recovered from zygotically but not from maternally expressed transcripts indicates that *d*NET-seq is specifically targeting the nascent transcriptome.

### *d*NET-seq reveals differences in transcription termination profiles between genes expressed in early and late embryos

Having confirmed that *d*NET-seq was capturing nascent RNA, we next investigated the distribution of Pol II density over transcript regions in early and late embryos. *d*NET-seq/S5P profiles (**Fig 3C-E and Supplemental Fig S3A-E**) do not show the characteristic higher read density near the promoter, as previously described in mammalian cells (Nojima et al. 2015; Mayer et al. 2015). This is most likely because Pol II typically pauses ~30-60 bp downstream of the transcription start site (TSS) (Kwak et al. 2013) and in our *d*NET-seq approach we enrich for RNAs longer than 60 nt (**Supplemental Fig S1F**); thus, we only record the position of polymerases that have transcribed at least 60 bp past the TSS.

We then turned our attention to the *d*NET-seq signal around the polyadenylation (pA) site. We found that in pre-MBT genes such as *snail* (**Fig 3D**), *tailless (tll)* (**Supplemental Fig S4A**) and *nullo* (**Supplemental Fig S4B**), *d*NET-seq/S5P signal ends soon after the pA site. In contrast, genes expressed in late embryos such as *Akap200* (**Fig 3E**), *His3.3A* (**Supplemental Fig S3H**) and *tsr* (**Supplemental Fig S3I**), have widespread *d*NET-seq/S5P signal on introns and regions downstream of the pA site. Analysis of RNA-seq datasets revealed that mRNAs encoded by these genes are efficiently spliced and 3’-end processed (**Supplemental Fig S3H, I**). Thus, *d*NET-seq/S5P is capturing newly synthesized transcripts that have not yet been spliced, as well as RNAs synthesized by Pol II complexes that continued to transcribe the DNA template after the mRNA was cleaved and polyadenylated at the pA site. We further noted that the 3’ UTR of genes with *d*NET-seq signal extending past the pA site is overlapped by another gene (*gurken, Nepl3,* and *IntS1,* respectively), which is transcribed in the opposite direction.

Next, we generated *d*NET-seq metaprofiles for pre-MBT and late genes. Metagene analysis of *d*NET-seq/S5P signal on pre-MBT genes expressed in early embryos confirmed that in this group of genes, *d*NET-seq signal is sharply reduced past the pA site (**Fig 3G**). The majority (65) of pre-MBT genes are isolated in the genome, with no other gene on either strand within a region of 500 bp downstream of the pA site, as shown for *tailless (tll)* (**Supplemental Fig S4A**). Another 26 pre-MBT genes are embedded in larger genes, as shown for *nullo*, which is located within a long intron of the *CG12541* gene (**Supplemental Fig S4B**). A smaller group of pre-MBT genes (21) have neighbouring genes located on either strand within a region of 500 bp downstream of the pA site, as shown for *Elba2* (**Supplemental Fig S4C**). We further identified 5 pre-MBT genes that have the 3’ UTR overlapped by an antisense convergent gene, as shown for *spook (spo)* (**Supplemental Fig S4D**). Notably, in these genes, the *d*NET-seq/S5P signal extended past the pA site (**Supplemental Fig S4C, D**).

To identify all the genes that are transcriptionally active in late embryos, we used a strategy adapted from GRO-seq analysis (Core et al. 2008) that relies on read density in gene desert regions as background reference for absence of transcription. Very large intergenic regions (gene deserts) were divided into 50kb windows, and read densities were calculated by dividing read counts in each window by the window length in bp (**Supplemental Fig S4E**). Genes with *d*NET-seq signal over the gene body (in RPKM) above the 90th percentile of read density for all intergenic regions analysed were considered to be transcriptionally active (**Supplemental Fig S4F**). We identified approximately 7 thousand active genes, with similar results obtained from *d*NET-seq/S5P and *d*NET-seq/S2P datasets (**Supplemental Fig S4G**). This set of active genes includes over 85% of the 3500 genes previously identified as actively transcribed after the mid-blastula transition (MBT) based on ChIP-seq experiments (Chen et al. 2013). Next, we divided the genes transcribed in late embryos into two groups, depending on whether their 3’ UTR was or was not overlapped by another convergent gene. The metagene analysis shows that when averaged across all transcribed genes, very low *d*NET-seq signal is detected at the pA site (**Fig 3H and Supplemental Fig S3G**), as expected assuming that when Pol II reaches this site, the nascent transcript is cleaved and polyadenylated and therefore there is no RNA left attached to the polymerase to be sequenced. However, in the case of late genes that have the 3’ UTR overlapped by an antisense (convergent) gene, the *d*NET-seq/S5P and *d*NET-seq/S2P signals increase again after the pA site (**Fig 3H and Supplemental Fig S3G**). Using the methodology described above to identify transcribed genes, we found that the vast majority (>80%) of overlapping convergent genes were transcriptionally active and only 219 genes were silent. The metagene analysis shown in **Fig 3I** clearly indicates that the detection of *d*NET-seq/S5P signal past the pA site is independent from transcription of the convergent overlapping gene.

In conclusion, the distribution profiles of *d*NET-seq/S5P and *d*NET-seq/S2P reads around the pA site suggest distinct patterns of transcription termination for genes that are either isolated in the genome or overlapped by another convergent gene. However, we cannot exclude the possibility that specifically in isolated genes, the lack of *d*NET-seq signal results from loss of CTD phosphorylation as Pol II transcribes past the pA site.

### Analysis of *d*NET-seq read density profiles

We next focused on the distribution of *d*NET-seq reads on exons and introns. The number of nascent RNA reads whose 3’ ends map at a particular genomic position is proportional to the number of Pol II molecules at that position. Thus, Pol II pause sites can be detected as local peaks in NET-seq read density (Churchman and Weissman 2011; Larson et al. 2014). However, because we performed *d*NET-seq with antibodies that recognize the Pol II CTD phosphorylated at either S5 or S2 positions, changes in read density may reflect dynamic phosphorylation of the CTD rather than Pol II pausing. We excluded signal resulting from splicing intermediates (i.e., reads that map to the very last nucleotide of introns and exons were discarded) in order to analyse only reads whose 3’ ends associate with Pol II active site. We analysed both S5P and S2P data. However, we were particularly interested in exploring the multiple peaks of dNET-seq/S5P signal observed on both exons and introns (**Fig 4A**), because co-transcriptional splicing has been linked to S5P in humans (Nojima et al. 2015).

**Figure 4.**
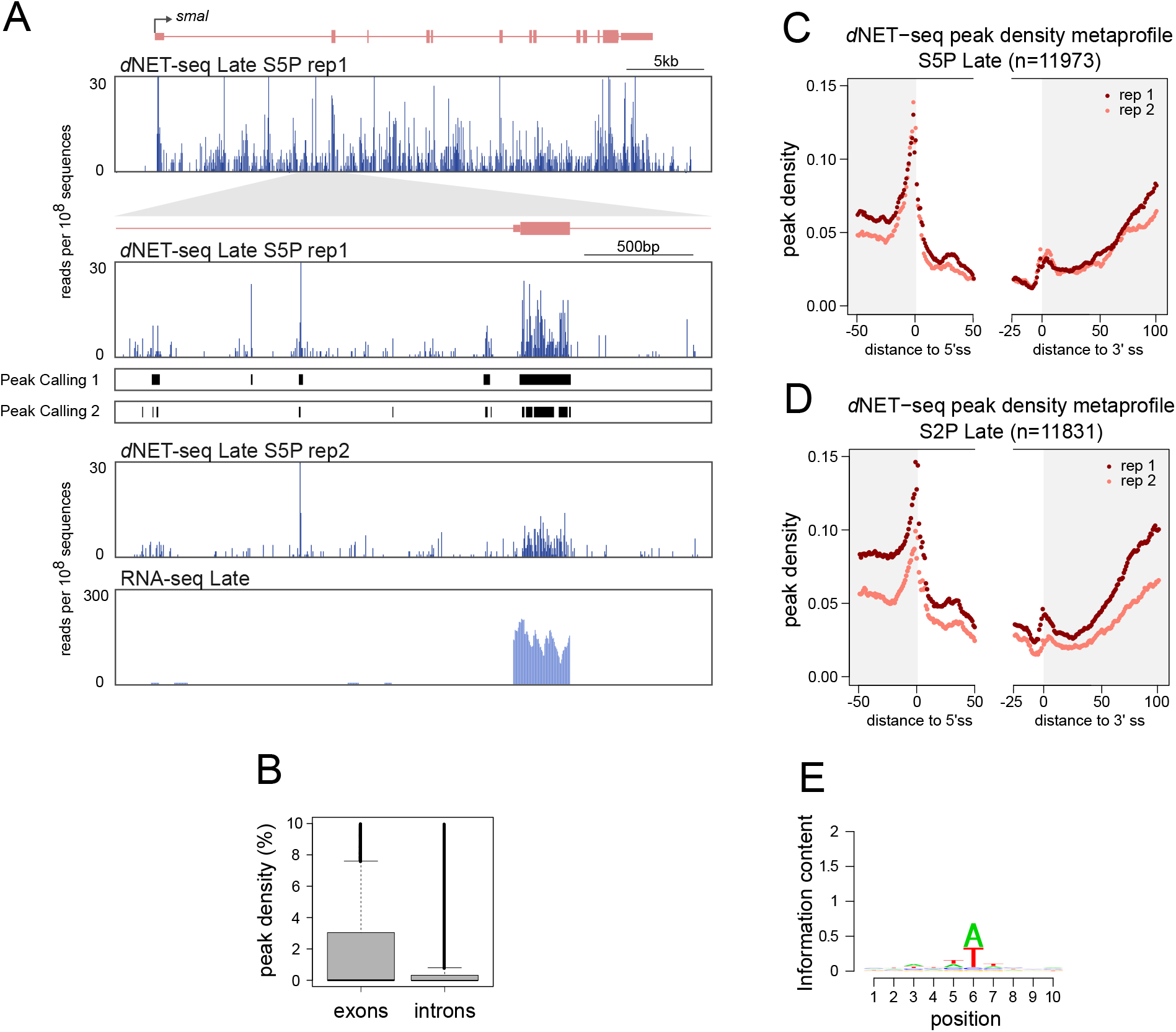
Analysis of *d*NET-seq read density profiles. (**A**) *d*NET-seq/S5P and RNA-seq profiles over the post-MBT gene *smoke alarm* (*smal*). The read number is depicted at two magnification levels in two biological replicates. For replicate 1, the line “Peak Caller 1” shows peaks called using the “large peaks” setting, which is appropriate for detecting larger regions of putative Pol II pausing. The line “Peak Caller 2” shows peaks called using the “small peaks” setting, which provides higher spatial resolution and has been used for subsequent analyses. RNA-seq data for the same regions is also shown. (**B**) Peak density in the exons and introns of transcriptionally active genes (*d*NET-seq/S5P, replicate 1). Peak density has been defined as the percentage of nucleotides within a given exon or intron that overlap with a significant peak. **(C-D)** Metagene analysis of peak density estimated from *d*NET-seq/S5P (**C**) and *d*NET-seq/S2P (**D**) data from late embryos. To calculate peak density for each position, we divided the number of introns that overlap with a peak at that position by the total number of introns. The last 50 nucleotides of exons, the first 50 nucleotides of introns, the last 25 nucleotides of introns, and the first 100 nucleotides of exons are shown. Only internal and fully coding exons from transcriptionally active genes that are at least 100 nucleotides long are shown. Exons shorter than 150 nucleotides contribute to both the exon end and start. Only introns that were at least 50 nt long were considered. **(E)** Sequence logo of nucleotide frequencies within a 10-nt window around the 5’ ends of NET-seq reads. The combined height of the bases at each position is proportional to the information content. Position 6 corresponds to the 5’-most nucleotide of the read. Putative internal priming reads, as well as reads mapping to the last nucleotide of exons or introns (possible splice intermediate and intron lariat reads) were ignored.

A difficulty when performing the systematic identification of *d*NET-seq peaks is that transcripts with higher initiation rates will contain more reads and thus peaks are more likely to be detected than in more lowly transcribed genes. To control for this confound, we developed a peak calling algorithm that detects regions where the local read density is significantly higher than expected by chance, given the over-all read density of the transcript (**Fig 4A;** see also the methods section). We emphasize that for detecting reads corresponding to splicing intermediates (**Fig 2D-F)**, we used a peak calling method that looks for significant single-nucleotide positions (Churchman and Weissman 2011), whereas this new method detects higher read regions of variable length.

We then aligned exons and introns on the splice sites and calculated the average peak density at each position. Exons have a higher over-all peak density than introns (mean proportion of nucleotides in peaks ~0.022/~0.015 for replicate 1/replicate 2 introns and ~0.043/~0.034 for replicate 1/replicate 2 exons; *P* < 2.2 * 10^−16^ for both replicates; two-tailed Mann-Whitney *U*-test with the peak densities of individual introns/exons as data points) (**Fig 4B**). This is consistent with previous reports indicating that Pol II elongation rate is decreased over exons in mammalian (Mayer et al. 2015; Jonkers et al. 2014) and *Drosophila* cells (Kwak et al. 2013). In addition, peak density is sharply increased around the 5’ splice site (**Fig 4C**). This could indicate either Pol II pausing or increased S5 phosphorylation associated with splice site recognition. We also cannot exclude that misaligned splicing intermediate reads are contributing to the observed increase in peak density around the 5’ splice site. The *d*NETseq/S5P profile around the 3’ splice site shows a higher peak density region just after the intron-exon boundary, and a progressive increase in average exonic peak density is further observed starting roughly 60 nt after the 3’ splice site (**Fig 4C**). A very similar peak density metaprofile is observed with the *d*NETseq/S2P dataset (**Fig 4D**).

A potential caveat of read density analysis is that nucleotide composition varies systematically across exons and introns. For example, exons tend to have a higher GC content than introns (Zhu et al. 2009). This could be problematic as NET-seq relies on MNase digestion of DNA and RNA to solubilize chromatin. MNase digestion of DNA is known to be sequence-biased, with most notably a preference for cleaving just 5’ of an adenine (Dingwall et al. 1981; Hörz and Altenburger 1981; Gaffney et al. 2012). An analysis of the 5’ ends of our reads revealed similar biases for MNase digestion of RNA (**Fig 4D**). This sequence preference could lead to artefactual variation in read density, with more reads being sampled from transcripts and transcript regions whose nucleotide composition is more similar to MNase digestion biases. To verify to what extent our results were affected by this confound, we performed a simulation to determine the expected distribution of reads based on the digestion bias alone (Supplementary Methods). We concluded that MNase biases are unlikely to explain either the enrichment of peaks in exons or the general profile of peak densities past the 3’ splice site.

### *d*NET-seq captures recursive splicing intermediates

Having shown that the spliceosome forms a complex with the elongating Pol II in *Drosophila* embryos, we asked when splicing takes place relative to transcription. We first looked at recursive splicing of long introns because this process involves the formation of inherently unstable intermediates that are more likely to be formed soon after the transcription of each intronic splice site (Pai et al. 2018). In recursive splicing, long introns are removed by sequential excision of adjacent sections involving separate splicing reactions, each producing a distinct lariat (Hatton et al. 1998). Recursively spliced intron segments are bounded at one or both ends by recursive sites or ratchet points (Burnette et al. 2005), which correspond to zero nucleotide exons consisting of juxtaposed 3’ and 5’ splice sites around a central AG|GT motif, where the vertical line represents the splice junction (**Fig 5A**).

**Figure 5.**
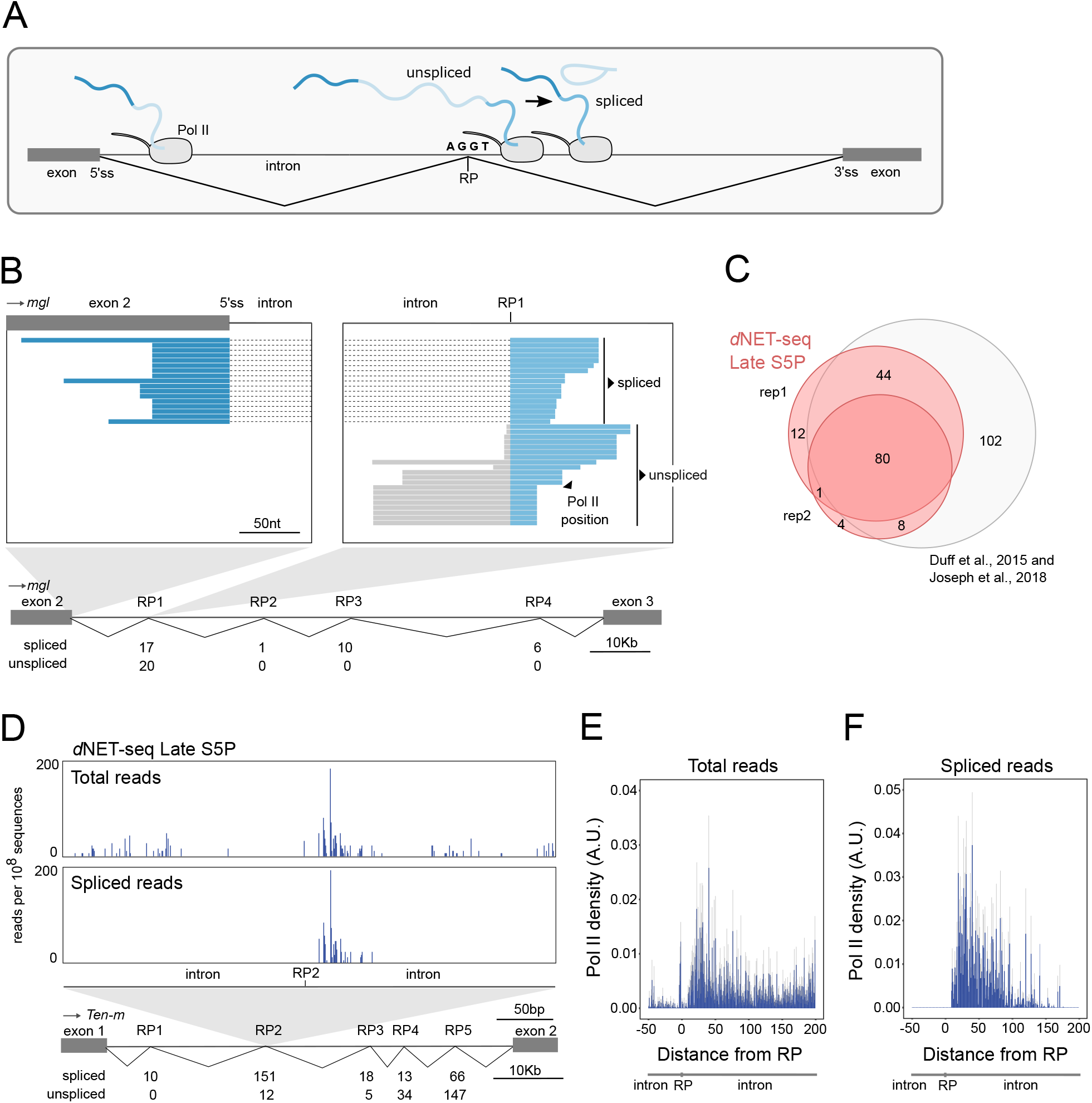
*d*NET-seq captures recursive splicing intermediates. **(A)** Schematic illustrating recursive splicing. A ratchet point (RP) with juxtaposed acceptor and donor splice site motifs is indicated. **(B)** Visualization of *d*NET-seq/S5P reads that align to the second intron of the *Megalin* (*mgl*) gene. Recursively spliced reads align to exon 2 (dark blue) and the intron after RP1 (light blue). Unspliced reads are depicted in grey. The number of spliced and unspliced reads at each RP in the intron is indicated. **(C)** Venn diagram comparing RPs identified in two *d*NET-seq/S5P biological replicates and in previously reported studies (Joseph et al. 2018; Duff et al. 2015). **(D)** Number of *d*NET-seq/S5P reads that have the 3’ end mapped around RP2 in the first intron of *Tenascin major* (*Ten-m*) gene. The top panel depicts all reads, and the bottom panel depicts only reads that have been spliced to RP1. **(E-F)** Meta-analysis with single nucleotide resolution of normalized *d*NET-seq/S5P reads around RPs (n=137) using all reads **(E)** or only reads spliced to the previous RP or exon **(F).**

To capture recursive splicing intermediates using *d*NET-seq, it is essential to have a good coverage of reads corresponding to nascent transcripts and spanning the splice junctions. The total number of reads resulting from nascent RNA in each *d*NET-seq dataset is depicted in **Supplemental Fig S1G**. By merging the sequencing information of overlapped paired-end reads (**Fig 1E**), we were able to sequence on average ~103 nucleotides per nascent RNA (**Supplemental Fig S1F and S1H**). Focusing on previously identified *Drosophila* ratchet points (Joseph et al. 2018; Duff et al. 2015), we found *d*NET-seq/S5P reads that span the junction between the canonical 5′ splice site at the end of the exon and the first ratchet point (RP1) internal to the downstream intron, as shown for the second intron of the *Megalin* gene (**Fig 5B**). Reads spanning the subsequent intronic RPs were also observed (**Fig 5B**). Overall, we detected *d*NET-seq/S5P and *d*NET-seq/S2P spliced reads supporting most of the previously identified recursive splicing events (**Fig 5C and Supplemental Fig S5A**).

Analysis of *d*NET-seq profiles around a RP reveals an enrichment of reads in a region located a few nucleotides downstream of the RP, as shown for RP2 in the first intron of the *Tenascin major* gene (**Fig 5D**). Noteworthy, most of these reads are already spliced to the previous RP (**Fig 5D**). A meta-analysis of *d*NET-seq/S5P reads around 137 RPs confirms that many spliced reads can be observed just downstream of RPs **(Fig 5E, F)**, demonstrating that splicing occurs soon after the transcription of intronic recursive sites. The observed enrichment of *d*NET-seq/S5P reads around RPs further suggests that recursive splicing and Pol II elongation rate may be kinetically coupled. In agreement with this view, a slow Pol II mutant enhanced recursive splicing of *Ubx* transcripts in *Drosophila* embryos (de la Mata et al. 2003).

### Splicing takes place as Pol II transcribes past the 3’ splice site

Having established that *d*NET-seq captures recursive splicing, we then asked whether *d*NET-seq reads spanning canonical exon-exon junctions were also detected **(Fig 6A)**. To identify splicing events, we considered all internal and fully-coding exons that are at least 100 nt long in actively transcribed genes in early and late embryos. For each splice junction, we counted how many reads had the 3’ end mapped to the first 100 nt of the exon. Only exons with at least 10 reads mapping to this region were considered (see the methods section for justification of the threshold). Then, the *d*NET-seq splicing ratio (SR) was calculated by dividing the number of spliced reads by the sum of the number of spliced and unspliced reads **(Fig 6A).** Reads could be counted as spliced or unspliced if their 3’ end mapped to within the first 100 nt of the exon and their 5’ end reached upstream of the 3’ splice site, allowing to check whether the intron was still present. A robust agreement of estimated SR values was observed between biological replicates for pre-MBT genes (**Supplemental Fig S6A**) and genes expressed in late embryos (**Fig 6B**).

SR values showed a bimodal distribution, with peaks at both extremes (SR = 0 and SR = 1; **Fig 6C** and **Supplemental Fig S6B**), indicating that a subset of junctions were always spliced immediately after transcription (SR=1), while others remained unspliced (SR=0). Notably, differences in SR distribution were found between *d*NET-seq/S5P and *d*NET-seq/S2P datasets (**Fig 6C, D and Supplemental Fig S6B, C**). Significantly lower SR values were detected for S2P compared to S5P (*P* < 2.2*10^−16^ from binomial regression, see Fig 6 legend for statistical details). In particular, junctions that were most frequently spliced (SR values close to 1) were predominantly captured by *d*NET-seq/S5P (**Fig 6C and Supplemental Fig S6B**). This clearly points to a preferential association between co-transcriptional splicing and S5 phosphorylation of Pol II CTD, as previously proposed in human cells (Nojima et al. 2015, 2018). We therefore continued to focus solely on the S5P datasets in the remainder of our splicing analysis.

**Figure 6.**
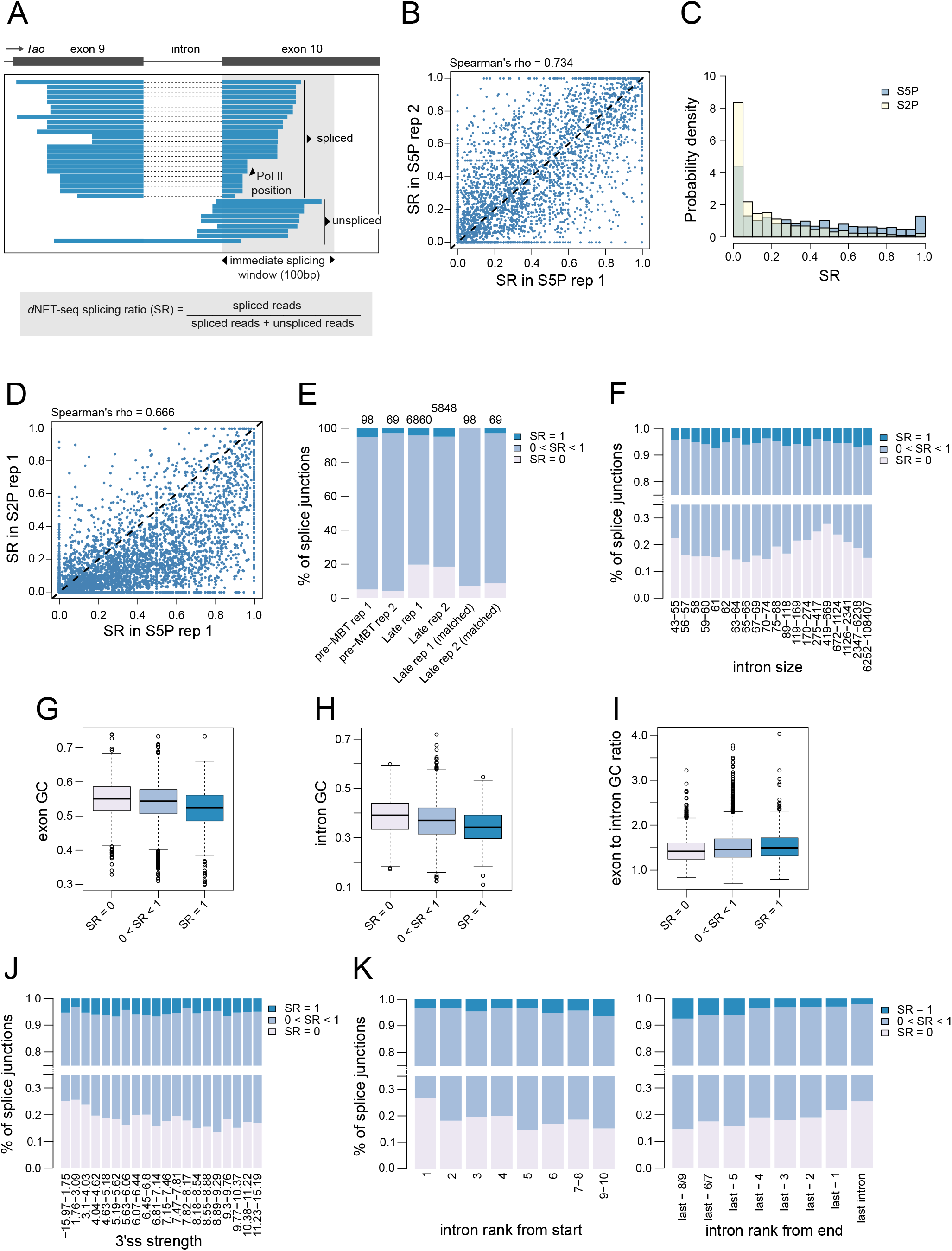
*d*NET-seq reveals immediate splicing at all intron sizes. **(A)** Visualization of *d*NET-seq/S5P reads that align to exon 10 of the *Tao* gene. For **(B-J)**, unless otherwise specified, only introns from transcriptionally active genes where the downstream exon is a fully coding internal exon at least 100 nt long were included. In addition, enough spliced/unspliced reads had to end within the first 100 exonic nucleotides that obtaining a splicing ratio (SR) of 0 or 1 by chance alone was highly unlikely (see the methods section for details). For genes expressed in late embryos, this threshold was 10 reads for both replicates. For pre-MBT genes, it was 14 for replicate 1 and 9 for replicate 2. For S2P data, we used a threshold of 10 to enable better comparison with S5P. **(B)** SR values estimated in two biological replicates of *d*NET-seq/S5P datasets from late embryos (Spearman correlation, ρ = ~0.734, *P* < 2.2 * 10^−16^; N = 3708). **(C)** Histogram of SR values for *d*NET-seq/S5P (N = 5626) and *d*NET-seq/S2P (N = 6888). To test the significance of the difference between S5P and S2P, a binomial regression with a logit link was performed without filtering by read number (N = 12833 for S5P; N = 13229 for S2P). The number of spliced and unspliced reads was specified as the dependent variable and the status of each data point as S5P or S2P was the sole predictor. The model predicted a splicing ratio of ~0.369/~0.394 for S5P replicate 1/2 and of 0.162/0.269 for S2P replicate 1/2. (**D**) SR values estimated in replicate 1 of *d*NET-seq/S5P and *d*NET-seq/S2P datasets from late embryos (Spearman correlation, ρ = ~0.666, *P* < 2.2 * 10-16; N = 4773). **(E)** Proportion of splice junctions in pre-MBT genes and genes expressed in late embryos classified according to their SR values. As many pre-MBT genes are single-intron, last introns were exceptionally included in this analysis. To make the two columns on the right, we used a subset of the genes expressed in late embryos (post-MBT genes) that was as similar as possible to the pre-MBT set in read number. Concretely, to match each pre-MBT gene, we picked the post-MBT gene that had the most similar total count of spliced and unspliced reads, making sure that every post-MBT gene only appeared in the subset once. **(F-K)** Several parameters of gene architecture show a relationship with SR. For the sample sizes and statistical tests used, see **Supplementary Table 1**. Note that in **(F, J-K)**, the bin ranges have been set so that intron numbers would be as equal as possible between bins.

We observed that only ~5% of splice junctions in pre-MBT genes were devoid of S5P reads spanning ligated exons and thus presented an SR of 0 (~5.10% replicate 1/~4.35% replicate 2), whereas in genes expressed in late embryos this proportion was ~20% (~19.72% replicate 1/~18.55% replicate 2) (**Fig 6E**; two-tailed binomial test for difference between late and pre-MBT, P ~ 6.141 * 10^−5^/9.44 * 10^−4^ (replicate 1/replicate 2)). However, this difference between pre-MBT genes and genes expressed in late embryos disappeared once the higher read density of pre-MBT genes had been controlled for (**Fig 6E**).

Thus, in most cases, *Drosophila* co-transcriptional splicing can occur when Pol II is still transcribing the downstream exon, implying an intron definition mechanism as previously proposed for *cerevisiae* (Carrillo Oesterreich et al. 2016). Consistent with this view, many *Drosophila* transcripts have relatively long exons separated by short introns – a gene architecture suggested to be conducive to intron definition. Very long introns flanked by short exons have instead been associated with exon definition (under which the downstream exon needs to be fully transcribed before splicing can take place) (Keren et al. 2010). A switch to exon definition was proposed once the size of the intron surpasses ~200 nt (Fox-Walsh et al. 2005). It is unclear, however, whether the choice between exon and intron definition is dependent on the absolute sizes of exons and introns, or rather the ratio of intron to exon size.

We found no relationship between the *d*NET-seq splicing ratio and exon length (**Supplemental Fig S6D, H** for replicates 1 and 2; see **Supplementary Table 1** for details on statistical significance and sample sizes for all of the gene architecture parameters discussed here). Regarding intron size, we found that introns with SR = 0 (and thus no evidence for intron definition) were, on average, indeed larger than other introns (**Fig 6F** and **Supplemental Fig S6G**). However, more careful examination revealed a more complex picture, with a lower proportion of introns with SR = 0 both for introns of intermediate size (~55-100 nt, which corresponds to ~55% of the introns studied) and for very large introns (>1000 nt) (**Fig 6F**). These intron sizes may thus be optimal for fast splicing. We also uncovered a lower exon to intron length ratio for introns with SR = 0 than for others (**Supplemental Fig S6E, I**; note that the difference was significant for replicate 2 and near-significant for replicate 1).

Taken together, these results suggest that although fast splicing (implying intron definition) is indeed skewed towards small introns flanked by large exons, there is also frequent and efficient intron definition for large introns. Our results are inconsistent with a threshold model, where splicing would systematically switch to exon definition after a given intron size is reached.

We also investigated the relationship between SR and several other gene architecture parameters. Firstly, as the GC content in exons and introns decreases, the proportion of introns with SR = 1 increases and the proportion with SR = 0 decreases, showing more efficient immediate splicing (**Fig 6G, H** and **Supplemental Fig S6K, L**). A similar effect is observed as the ratio of the downstream exon GC content to intron GC content increases (**Fig 6I** and **Supplemental Fig S6M**). Thus, the most efficient immediate splicing is observed when the GC content is low in both exons and introns but higher in exons than introns. Secondly, as 3’ splice site strength increases, the proportion of introns with SR = 0 decreases (**Fig 6J** and **Supplemental Fig S6N**). We found no significant relationship between SR and 5’ splice site strength (**Supplemental Fig S6F, J**). However, we replicated previous observations (Khodor et al. 2012) that transcripts with only a single intron tend to be spliced less efficiently than multi-intron ones, with a higher proportion of SR = 0 introns (**Supplemental Fig S6O, P**). Thirdly, similarly to previous reports (Khodor et al. 2011, 2012; Herzel et al. 2018), we uncovered an effect of exon rank, whereby exons that are more central appear to be spliced more efficiently (**Fig 6K** and **Supplemental Fig S6Q**).

In conclusion, *d*NET-seq reveals that splicing can occur immediately as Pol II transcribes past the 3’ splice site, yet many nascent transcripts remain unspliced. As expected, we found a relationship between the proportion of spliced reads and intron size. However, immediate splicing was observed at all intron sizes.

### Immediate splicing associates with specific read density profiles

Analysis of individual *d*NET-seq/S5P profiles around constitutive splice junctions with a high splicing ratio frequently revealed an enrichment of reads downstream of the 3’ splice site, coincident with the appearance of spliced reads, as shown for the *cno* gene (**Fig 7A**). In contrast, profiles around a constitutive splice junction with SR = 0 had an accumulation of reads further along the exon, as shown for the *ND-51* gene (**Fig 7B**). A clearly distinct type of profile was observed on skipped exons, on which very few reads were observed, as shown for the *zip* gene (**Fig 7C**).

**Figure 7.**
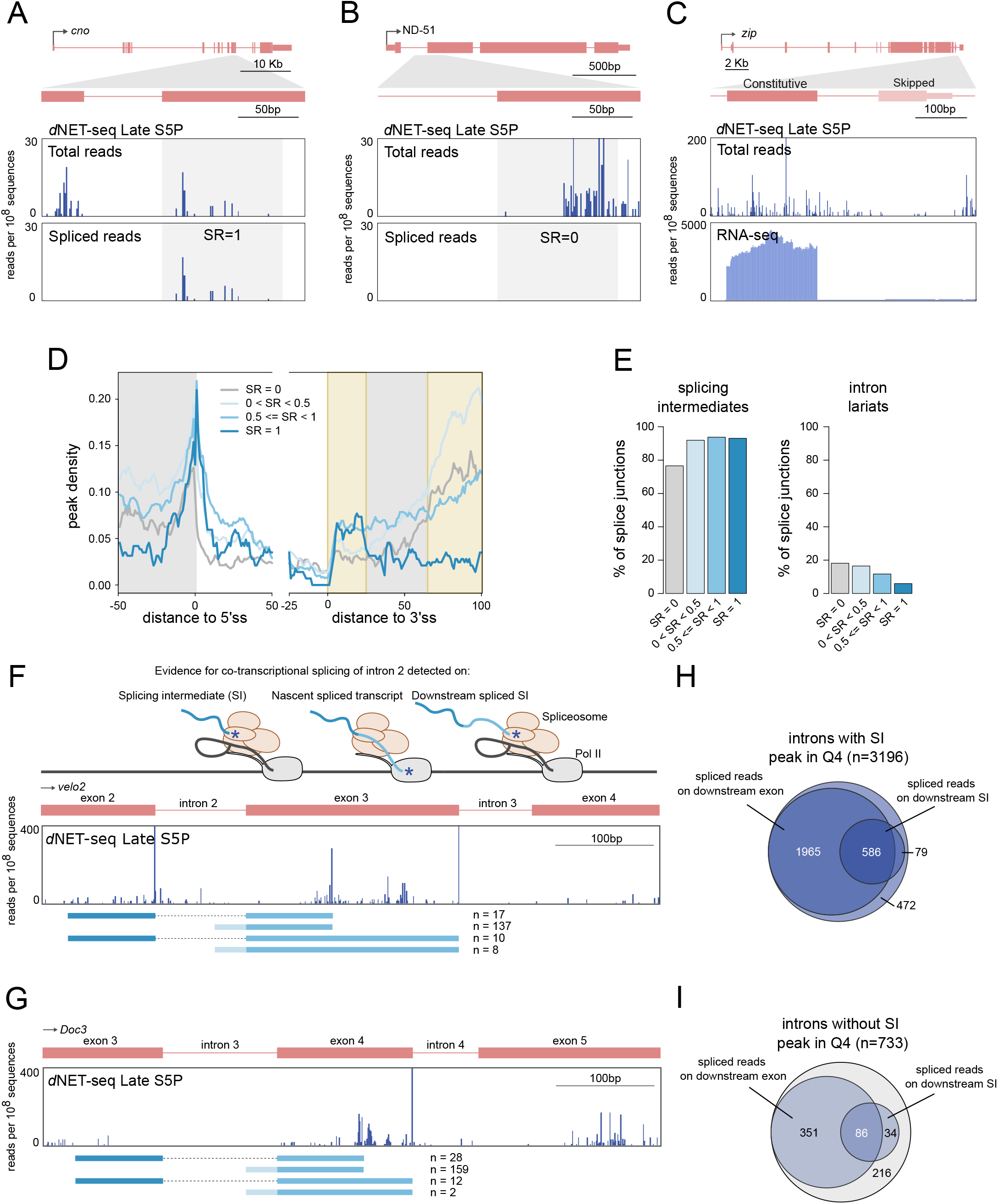
Immediate splicing associates with higher density of *d*NET-seq signal. **(A-C)** *d*NET-seq/S5P profiles surrounding the indicated exons in the post-MBT genes *cno* (**A**), *ND-51* (**B**), and *zip* (**C**). The top panels depict all reads. The bottom panels depict either the 3’ end coordinate of reads that span the splice junction **(A, B)**, or the RNA-seq profile (**C**). **(D)** Metagene analysis of peak density estimated from *d*NET-seq/S5P datasets from late embryos (replicate 1) for different ranges of SR values. To calculate peak density for each position, we divided the number of introns that overlap with a peak at that position by the total number of introns. The last 50 nucleotides of exons, the first 50 nucleotides of introns, the last 25 nucleotides of introns, and the first 100 nucleotides of exons are shown. Only internal and fully coding exons from transcriptionally active genes that are at least 100 nucleotides long are shown (N = 4783). In addition, at least 10 spliced/unspliced reads had to end within the first 100 nucleotides of the exon. (**E**) The proportion of introns with at least one read whose 3’ end maps to the final position of the upstream exon (putative splicing intermediates) or to the final position of the intron (putative intron lariats) in *d*NET-seq/S5P late replicate 1. (**F, G**) *d*NET-seq/S5P profiles on the indicated regions of the *velo2* and *Doc3* genes. Below, spliced reads are depicted. Asterisks denote 3’ OH ends. **(H, I)** Venn diagrams showing how many junctions with or without a splicing intermediate peak are covered by spliced reads or have a downstream splicing intermediate covered by spliced reads.

We next performed a meta-analysis of *d*NET-seq/S5P peak densities over different exonic and intronic regions for introns with differing SR values. A difficulty of analysing the peak density profile around the 3’ splice site is that reads mapping to the final nucleotide of the intron may represent intron lariats and thus be non-nascent. We removed reads mapping to this position prior to peak calling. However, through misalignment, intron lariat reads may also map to the few nucleotides around the 3’ splice site and thus still affect the final meta-profile. In order to minimize the impact of such misalignment events, we excluded from the meta-analysis all introns with a read mapping to the final intronic nucleotide, as this is expected to also discard the introns most likely to contain misaligned intron lariat reads (**Fig 7D, Supplemental Fig S7A**). When such filtering is not done, a sharp peak is observed around the 3’ splice site, notably when SR values are low (**Supplemental Fig S7 B-C**). It appears that intron lariat reads are primarily captured for introns with low splicing ratios (**Fig 7E and Supplemental Fig S7D**). This contrasts with splicing intermediate reads from the end of the upstream exon, which are associated to higher splicing ratios instead (**Fig 7E and Supplemental Fig S7D**).

The peak density profile downstream of the 3’ splice site contains two primary regions of interest (highlighted in **Fig 7D and Supplemental Fig S7 A-C, G-H**). The first is a region of increased peak density located roughly 10-20 nucleotides after the 3’ splice site. This peak is independent of the presence of putative intron lariat reads (**Fig 7D, Supplemental Fig S7 A-C**), and is more prominent when the SR is higher (**Fig 7D, Supplemental Fig S7 A-C, G-H**).

The second feature of interest is a rapid increase in peak density ~60 nucleotides into the exon. Contrary to the first region of interest, the peak density in this region is higher when SR is *lower* (**Fig 7D, Supplemental Fig S7 A-C, G-H**). We found no evidence that this increase in peak density could be a result of MNase digestion biases (**Supplementary Methods**). As discussed above, this pattern of increased *d*NET-seq/S5P density could result from either a local enrichment of CTD S5 phosphorylation or from an accumulation of Pol II due to a decreased elongation rate. In the latter case, one possible explanation for such a slow-down over exonic regions could be their relatively higher GC content when compared to intronic sequences. The peak is indeed more prominent in introns where the ratio of exonic to intronic GC content is higher (**Supplemental Fig S7 E-F**). However, when SR=1, then even exons where this ratio is high do not display the peak (**Supplemental Fig S7 E-F**). Moreover, introns with higher SR tend to have *higher* exonic to intronic GC ratios (**Fig 6I, Supplemental Fig S6M**). Hence, the co-variation with SR cannot be explained simply through the correlation between SR and GC content patterns. This suggests that there is an effect of the SR beyond any GC content effect. Potentially, this region could correspond to where delayed splicing starts (because these events have low immediate splicing ratios). Note that we obtained qualitatively similar meta-profiles when considering read rather than peak densities (**Supplemental Fig S7 G-H**). Our results therefore cannot be explained through any biases introduced by the peak calling approach.

Finally, we asked how prevalent co-transcriptional splicing is in the developing *Drosophila* embryo. As shown in **Fig 2E**, approximately 80% of exons in pre-MBT genes and genes expressed in late embryos covered by a high density of NET-seq reads have a peak corresponding to the splicing intermediate formed after cleavage at the 5′ splice site but before exon-exon ligation. The majority of these exons (~85%) were covered by *d*NET-seq reads that span the junction to the downstream exon either directly on nascent transcripts or indirectly on splicing intermediates formed by cleavage at the 5’ splice site of the downstream exon **(Fig 7F, H)**, confirming that they are co-transcriptionally spliced. We then focused on those exons for which no splicing intermediate spike was detected by the peak calling algorithm, as illustrated for the *Doc3* gene **(Fig 7G)**. Over 71% of these exons were also covered by *d*NET-seq reads that span the junction to the downstream exon either directly on nascent transcripts or indirectly on splicing intermediates formed by cleavage at the 5’ splice site of the downstream exon **(Fig 7I)**, arguing that even exons without a detectable splicing intermediate peak are co-transcriptionally spliced. Altogether, our *d*NET-seq results support co-transcriptional splicing for over 95% of the analysed exons.

## DISCUSSION

In this study, we used NET-seq to map Pol II with the CTD phosphorylated on either S5 or S2 positions over the bodies of genes that become transcriptionally active in *Drosophila* embryos during the initial stages of development. The use of embryos allowed us to perform an important test to verify that the captured RNA is truly nascent. Indeed, early *Drosophila* embryos contain abundant mRNAs that are transcribed during oogenesis and loaded into the egg. In our analysis of early embryos, these maternal transcripts were readily detected by RNA-seq, but not by *d*NET-seq.

Transcription in the *Drosophila* early embryo begins in mitotic cycle 8 for a few genes (Erickson and Cline 1993; Pritchard and Schubiger 1996) and then the number of active genes gradually increases until cycle 14 (Kwasnieski et al. 2019). Notably, the initial mitotic cycles have a duration of approximately 10 minutes, with cycle 13 taking about 21 minutes and cycle 14 lasting for at least 65 minutes (Ji et al. 2004). Thus, assuming an elongation rate of 2.4-3.0 kb per minute (Fukaya et al. 2017), there are significant constraints on the transcription of genes that are active before the end of cycle 13 (pre-MBT genes). Although pre-MBT genes are on average shorter than genes expressed at later stages of development (Artieri and Fraser 2014; Hoskins et al. 2011), longer genes are nevertheless transcribed before MBT. However, before cycle 14, transcriptional elongation of long genes is either prematurely terminated (Sandler et al. 2018) or aborted (Shermoen and O’Farrell 1991; Kwasnieski et al. 2019). It has also been reported that pre-MBT expression is associated with the generation of DNA damage due to stalling of DNA replication at transcriptionally engaged loci (Blythe and Wieschaus 2015).

Our *d*NET-seq analysis revealed a novel feature of pre-MBT genes that most likely contributes to reduce conflicts between transcription and replication during the short interphases of early embryos. We found that in the majority of pre-MBT genes *d*NET-seq/S5P signal is not detected beyond the pA site, suggesting that Pol II terminates soon after cleavage and polyadenylation. We also observed that these pre-MBT genes tend to be isolated from other transcriptional units. However, a small subset of pre-MBT genes have neighbouring genes and in those cases *d*NET-seq/S5P signal extends past the pA site. In contrast to pre-MBT genes, many of the genes expressed after MBT have the 3’UTR overlapped by another gene transcribed in the opposite direction and show high *d*NET-seq/S5P and S2P signal past the pA site. The finding that very low *d*NET-seq/S5P and S2P signal is detected at the pA site and increases sharply thereafter suggests that nascent transcripts are efficiently cleaved and polyadenylated, yet Pol II remains associated with the DNA template synthesizing RNA past the pA site. Indeed, analysis of RNA-seq datasets confirmed that mRNAs transcribed from these late genes are normally 3’-end processed. Altogether, these observations suggest that transcription termination of isolated genes occurs soon after cleavage and polyadenylation, whereas in genes with overlapping neighbours Pol II continues to transcribe into the gene 3’-flanking region after passage of the pA site. Notably, the proportion of isolated genes is significantly higher among pre-MBT genes (65/117 or ~56%) than among late genes (1251/7233 or ~17% of all transcriptionally active genes for replicate 1 S5P) (*P* < 2.2 * 10^−16^, one-tailed binomial test). Possibly, pre-MBT genes in the *Drosophila* genome may be under selection to avoid overlaps with other genes, thereby minimizing potential transcriptional interference problems such as collisions involving DNA polymerase complexes or Pol II transcribing opposite template strands (Proudfoot 2016).

Our observations further reveal an unexpected link between delayed transcriptional termination and the presence of an overlapping convergent gene in the *Drosophila* genome. It is intriguing that Pol II persists transcribing after cleavage and polyadenylation of the nascent mRNA, specifically when there is a convergent overlapping gene and regardless of its transcriptional status. One possibility is the induction of local conformational changes in chromatin and/or Pol II. Indeed, pioneer transcription factors can potentially regulate gene expression at later stages of development by inducing significant chromatin conformational changes during early embryonic development (Blythe and Wieschaus 2016; Schulz et al. 2015). Although such chromatin remodelling events can potentially influence the interaction of Pol II with DNA downstream of the pA site, further studies are needed to understand how transcription termination is regulated during early *Drosophila* development.

A classical view in the splicing field is that for some introns, the 5’ and 3’ splice sites are recognized directly by ‘intron definition’. For other introns, spliceosome assembly starts with the recognition of the downstream exon, and only at a later stage a cross-intron complex is formed (’exon definition’) (Robberson et al. 1990; Berget 1995; Sterner et al. 1996; Fox-Walsh et al. 2005). These alternative models are supported by *in vitro* evidence, as well as by *in vivo* reporter gene experiments (Robberson et al. 1990; Berget 1995; Fox-Walsh et al. 2005; De Conti et al. 2013). There is also indirect evidence, for example from analysis of exon-intron size patterns (Berget 1995). However, these models have never been directly tested *in vivo* at a transcriptome-wide level. It thus remains uncertain how intron definition and exon definition operate *in vivo* and what determines the choice between them.

Introns that show splicing completion before Pol II has reached the end of the downstream exon are necessarily intron-defined, as exon definition requires the presence of the 5’ splice site of the downstream intron. Therefore, introns with an SR above 0 represent an experimentally-determined set of introns that are spliced *via* intron definition at least part of the time, allowing us to test long-held assumptions in the field. Notably, *in vitro* and reporter gene work suggested that *Drosophila* uses intron definition for introns smaller than ~200-250 nt and exon definition for larger introns (Fox-Walsh et al. 2005). Other studies have suggested that the crucial factor is not just intron size but the relative size of the intron compared to the flanking exons (Sterner et al. 1996).

Using *d*NET-seq, we have found that although there is a relationship between intron size and the proportion of introns with SR > 0, intron definition can be observed at all intron sizes. Our results are thus inconsistent with a simple “threshold” model, where splicing would systematically switch to exon definition at a particular intron size. Rather, we observe an increased proportion of introns with SR > 0 within an optimal intron size range of ~55-100 nt. The median intron size in our data set is 71 nt. It thus appears that intron definition is best optimized for introns of a “typical” size, and is less efficient when introns are unusually small or large. Intriguingly, however, for very large introns (>1000 nt), the proportion with SR > 0 increases again, suggesting that specific mechanisms may have evolved for the intron-defined splicing of very large introns. The tendency for faster splicing kinetics with average intron sizes has also been reported using metabolic labelling in both *Drosophila* (Pai et al. 2017) and human cells (Windhager et al. 2012).

We propose that the tethering of the upstream exon to Pol II (de Almeida and Carmo-Fonseca 2008) could be crucial for the intron-defined splicing of such large introns. Indeed, introns with SR > 0 where the upstream exon has one or more putative splicing intermediate reads tend to be larger than ones without splicing intermediate reads (one-tailed Mann-Whitney *U*-test, W = 779623, P ~ 0.003, N = 4589; exon size and read number filtering has been applied like for the gene architecture analysis).

In addition, Pai et al. (2017) found that GC poorer introns were spliced faster. We uncovered a similar relationship with GC content, with the highest proportion of introns with SR > 0 observed when both the exon and the intron were AT-rich but the exon had a higher GC content than the intron. We emphasize that the gene architecture parameters that we have investigated are not independent of each-other. For instance, exon GC content is correlated negatively with exon rank from start (Spearman’s ρ ~ −0.098; P ~ 1.196 * 10^−12^). Further work is thus needed to distinguish between causative and merely correlative factors. In addition, it must be determined how the GC content effect relates to patterns of nucleosome density (Gelfman et al. 2013).

Our results indicate that the majority (>80%) of *Drosophila* introns expressed in late embryos can be spliced through intron definition. A limitation of our work is that introns that show no evidence for splicing completion during transcription of the downstream exon could still be spliced *via* intron definition, if their splicing takes too long for it to be detected during the 100 nt window studied. Therefore, we do not know whether introns with a SR of 0 are exon-defined, intron-defined but spliced when Pol II has elongated past the first 100 exonic nucleotides, or a mixture of both. Similarly, introns with a splicing ratio between 0 and 1 could either always be intron-defined, or they could use either exon or intron definition depending on the splicing event.

Taken together, our results show that splicing in *Drosophila* embryos can be completed soon after transcription of the 3’ splice site, as previously reported in other cellular models (Carrillo Oesterreich et al. 2016; Reimer et al. 2021; Martin et al. 2013; Sousa-Luís et al. 2021). However, a question that remains open is whether splicing influences the kinetics of Pol II elongation. A splicing-dependent Pol II pausing near the 3’ splice site was first suggested by the Beggs group (Alexander et al. 2010; Chathoth et al. 2014), and evidence for Pol II pausing at exon boundaries was later detected in mammalian cells using NET-seq (Nojima et al. 2015; Mayer et al. 2015). In contrast, a recent PRO-seq analysis found no splicing-associated Pol II pausing (Reimer et al. 2021). A splicing-related accumulation of S5P Pol II along gene bodies was also observed using chromatin immunoprecipitation in HeLa cells (Batsché et al. 2006) and in yeast (Alexander et al. 2010; Chathoth et al. 2014). However, it remains unclear whether a local increase in NET-seq read density results from slower Pol II elongation or represents a region of preferential CTD Ser5 phosphorylation, with no concomitant Pol II pausing. In this regard, splicing-associated higher NET-seq density profiles were observed with S5P antibodies but not with antibodies to unphosphorylated CTD (Sheridan et al. 2019; Nojima et al. 2015). Clearly, future studies are needed to assess the distribution of nascent transcripts independently from the Pol II phosphorylation status.

Irrespective of Pol II pausing, the results of our *d*NET-seq/S5P read density analysis around canonical and recursive splice sites strongly suggest that CTD serine 5 can be dynamically re-phosphorylated along gene bodies in a splicing-dependent manner, as previously proposed (Nojima et al. 2018; Harlen and Churchman 2017). Moreover, our analysis of nascent *Drosophila* transcripts associated with Pol II S5P CTD revealed highly heterogeneous splicing dynamics. Indeed, while only a minority of splice junctions were devoid of reads spanning ligated exons (corresponding to an SR of 0), many introns remained unspliced in our analysis. In agreement with these observations, a recent single-molecule study of nascent RNA dynamics in live cells found large kinetic variation in intron removal for single introns in single cells (Wan et al. 2021). Is it relevant for cells whether any particular intron is rapidly excised after Pol II transcribes the 3’ splice site or is left unspliced while the transcript elongates? Is the decision to be rapidly excised or left unspliced dependent on a stochastic process? What are the consequences for gene expression of fast versus delayed splicing? Addressing these questions will likely dependent on further advances in methodologies to study splicing in time and space at a transcriptome-wide level.

## METHODS

### Embryo collection

*Drosophila melanogaster* flies (Oregon R (OrR) strain) were raised at 25°C, in polypropylene vials containing standard enriched culture medium (cornmeal, molasses, yeast, soya flour, and beetroot syrup). Three-day-old flies (counting from pupae eclosion) were fattened in culture medium supplemented with fresh yeast for 2 days. Embryos were collected into apple juice-agar plates supplemented with fresh yeast using appropriate cages containing approximately 200 flies each. To avoid female retention of older embryos, three pre-collections of 30min each were made before the first collection of embryos. To maximize egg laying, and avoid larvae contamination, adult flies were transferred to clean embryo collection cages every day over five days. For early stage embryos (2-3 hours after egg-laying), adult females were allowed to lay eggs for 1 hour in apple juice-agar plates. Plates were subsequently collected and embryos were aged at 25°C for 90 min. During the following 30 min, embryos were harvested from the plates, dechorionated in 50% bleach solution for 2 min, and washed once in Phosphate Buffered Saline supplemented with 0.1% Tween-20 (PBT) and twice in deionized water. In order to discard older embryos (stage 6 and older), manual staging of collected embryos was performed with the help of forceps and under a magnifier scope. Embryos were then resuspended in a solution containing 120mM NaCl and 0.04% Triton, and washed twice with 120mM NaCl solution. At the end of the 3 hours collection, the solution was removed and embryos were frozen in liquid nitrogen and stored at −80°C. For the late stage (4-6 hours), eggs were laid for 2 hours and aged at 25°C for 3.5 hours. Embryo collection and processing was similar to the early stage embryos, but in this case, no manual staging was performed.

### Embryo DNA staining

For each embryo collection, and after dechorionation, a representative embryo sample was collected from the total pool and fixed in a scintillation flask, using a solution containing 1 volume of 4% formaldehyde in PBT and 4 volumes of heptane, for 20 min at 100rpm. The lower aqueous phase solution was subsequently removed, 4ml of methanol was added and embryos were shaken vigorously during 1 min. Embryos were then collected from the bottom of the scintillation flask, washed twice with methanol, and frozen at −20°C in methanol. To rehydrate the embryos, they were washed for 5 min each, with 3:1, 1:1, and 1:3 mix solutions of methanol:PBT. Embryos were subsequently washed twice in PBT and incubated with 1:5000 Sytox green (Invitrogen), supplemented with 5μg/ml RNase A (Sigma-Aldrich) in PBT for 15 min. After washing with PBT, embryos were mounted in fluorescence mounting medium (Dako) and examined in a Zeiss AxioZoom V16 Fluorescence Stereo Microscope for image acquisition and embryo staging. Images were processed using ImageJ software (NIH).

### *d*NET-seq and library preparation

The *d*NET-seq protocol was adapted from mNET-seq (Nojima et al. 2016). Briefly, 300 ul of frozen embryos was resuspended in 3.5ml of Buffer B1 (15mM HEPES-KOH, pH 7.6; 10mM KCl; 5mM MgCl2; 1mM DTT; 0.1mM EDTA; 0.35M Sucrose; 4μg/ml Pepstatin; 10mM Sodium Metabisulfite; 0.5mM EGTA supplemented with complete EDTA free protease inhibitor (Roche) and PhoSTOP (Roche)). Embryos were homogenised in a Dounce homogenizer with 11x strokes using a tight pestle on ice. The suspension was centrifuged at 7700g for 15 min at 4°C, the supernatant was discarded and the white pellet containing the nuclei was resuspended in 500 μl of Buffer B1. The suspension was again homogenised in the Dounce with 4x strokes and loaded without mixing on the top of buffer B2 (15mM HEPES-KOH, pH 7.6; 10mM KCl; 5mM MgCl2; 1mM DTT; 0.1mM EDTA; 0.8M Sucrose; 4μg/ml Pepstatin; 10mM Sodium Metabisulfite; 0.5mM EGTA supplemented with complete EDTA free protease inhibitor (Roche) and PhosSTOP (Roche)). The suspension was centrifuged at 1310g during 30 min at 4°C, and the pellet was resuspended with 125 μl of NUN1 buffer (20mM Tris-HCl (pH 7.9); 75 mM NaCl; 0.5mM EDTA and 50% Glycerol). 1.2ml of Buffer NUN2 (300mM NaCl, 7.5mM MgCl2, 1% NP-40, 1M Urea supplemented with complete EDTA free protease inhibitor (Roche) and PhoSTOP (Roche) was mixed with the nuclei and incubated on ice during 15 min preforming short vortex every 3 min. Chromatin was then centrifuged at 10000g for 10 min at 4°C, washed with 100 μl of 1x MNase Buffer and incubated in 100μl of MNase reaction mix (1x MNase buffer and 30 gel unit/μl MNase (New England Biolabs)) during 3 min at 37°C with mix at 1400rpm. The reaction was stopped with 10 μl of 250mM EGTA, centrifuged at 10000g for 5 min at 4°C, and the supernatant containing the solubilized chromatin was recovered. For the early embryos sample, 2x 300μl of embryos were prepared in parallel and pooled together after the chromatin solubilization. Immunoprecipitation of Pol II-RNA complexes was performed using 50μl of Protein G Dynabeads (Thermo Fisher Scientific), pre-incubated over night with 5μg of the correspondent antibody: anti-Pol II CTD S5P (ab5131 Abcam) or anti-Pol II CTD S2P (ab5095 Abcam) in 100μl NET2 (50mM Tris-HCl pH 7.4; 150mM NaCl and 0.05% NP-40) and washed 3 times with NET2. Beads were incubated with the solubilized chromatin in 1ml total volume of NET2 during 1 hour at 4°C, washed 7 times with 500μl of NET2 and once with 100μl of PNKT (1x PNK buffer and 0.1% tween) before incubation during 6 min in 50μl of PNK reaction mix (1x PNKT, 1 mM ATP and 0.05 U/ml T4 PNK 3′phosphatase minus (NEB) in a thermomixer at 37°C and 1400rpm. After washing the beads with NET2, long RNA fragments were isolated using Quick-RNA MicroPrep (Zymo research): 300μl of RNA Lysis Buffer in 33% EtOH was mixed to the beads by pipetting. Beads were discarded and the suspension was loaded into a Zymo spin column that was centrifuged at 10000g during 30sec. The column was washed once with 400μl RNA prep buffer and twice with 700μl and 400μl RNA wash buffer respectively. RNA was then eluted in 15μl of DNase/RNase-Free water (zymo) and stored at −80°C. 100ng of RNA was used to prepare each library, following the standard protocol of the Truseq small RNA library prep kit (Illumina). After adapter ligation and reverse transcription, the libraries were PCR amplified using 16 PCR cycles and cDNA libraries were fractionated in the gel between 130 to 300bp. The libraries were sequenced using PE-150 on the Illumina HiSeq X platform by Novogene Co., Ltd.

## QUANTIFICATION AND STATISTICAL ANALYSIS

### *d*NET-seq data processing

Adapter sequences were removed from all *d*NET-seq paired-end samples using Cutadapt (version 1.18) (Martin 2011) with the following parameters: -a TGGAATTCTCGGGTGCCAAGG -A GATCGTCGGACTGTAGAACTCTGAAC -m 10 -e 0.05 --match-read-wildcards -n 1. Paired-end read merging was performed using bbmerge.sh from BBMap (Bushnell et al. 2017) with the ‘xloose’ parameter. Merged reads were then aligned to the *Drosophila* reference genome (*dm6*; Ensembl release 95) (Cunningham et al. 2019) using STAR (version 2.6.0b) (Dobin et al. 2013) with –chimSegmentMin set to 20. Only uniquely mapped reads were considered, extracted using SAMtools (version 1.7) (Li et al. 2009) with -q set to 255. Exceptionally, in **Supplemental Fig S2A**, HiSat2 was used, using the same dm6 genome annotation file for genome indexing and using default parameters, with the --no-discordant –no-mixed flags set. PCR internal priming events generated during library preparation were removed using a custom Python script (Prudêncio et al. 2020) with the following parameters: -a TGG.. -s paired. To obtain single-nucleotide resolution, a custom Python script (Prudêncio et al. 2020) was used to extract the 5′ end nucleotide of read 2 (after trimming) in each sequencing pair, with the directionality indicated by read 1 (**Fig 1E**).

### Publicly available RNA-seq datasets used

Publicly available *Drosophila* embryonic transcriptome sequencing data (Poly(A) RNA-seq), performed in developmental stages similar to the dNET-seq early and late samples, were used in this study. RNA-seq datasets corresponding to cycle 14B (Lott et al. 2011) were obtained from the Gene Expression Omnibus (GEO) (samples GSM618409, GSM618410, GSM618421 and GSM618422 from dataset GSE25180). RNA-seq datasets from 4-6h old embryos were obtained from modENCODE project PRJNA75285 (Graveley et al. 2011) (accessions SRR023696, SRR023746, SRR023836, SRR035220, SRR023669, SRR035405, SRR035406, SRR024014 and SRR023539).

### RNA-seq data processing

Adapters were removed from all datasets using Trim Galore (version 0.4.4) (http://www.bioinformatics.babraham.ac.uk/projects/trim_galore/; last accessed 26 April 2020). Datasets from modENCODE and GEO were aligned to the *dm6 Drosophila* reference genome (Ensembl release 95) (Cunningham et al. 2019) using STAR (version 2.6.0b) (Dobin et al. 2013) with -- chimSegmentMin set to 20. Stringtie (version 1.3.3b) (Pertea et al. 2015) was used to quantify normalized gene expression as Transcripts Per Kilobase Million (TPM) values with the following parameters: -a 5 -e. In addition, the isoform list was provided together with the -G parameter corresponding to the *dm6* (Ensembl release 95) GTF file. Genes with TPM values above 2 were considered to be expressed.

### Selection of genes and isoforms for analysis

Similar to a previous GRO-seq analysis (Core et al. 2008), we used the read density of very large intergenic regions (gene desert regions) to define the reference for absence of transcription. Gene deserts were divided into 50kb windows, and *d*NET-seq read densities were calculated by dividing the read counts in each window by the window length (in bp). Read counts per window were obtained with bedtools genome coverage (version 2.27.1-1-gb87c465) (Quinlan and Hall 2010), and an arbitrary density threshold was defined as the 90th percentile of the read density distribution (**Supplemental Fig S4A**). Transcripts whose gene body *d*NETseq read density exceeded this threshold were considered to be transcriptionally active (**Supplemental Fig S4B**). For all of the analyses performed on late genes, only transcriptionally active genes were considered. In addition, only one isoform per gene was considered for all analysis, selected as the isoform with the highest RPKM value in the RNA-seq dataset for the corresponding developmental stage.

The coordinates of previously identified pre-MBT genes were obtained from Chen et al. (2013) and converted to *dm6* coordinates using the *liftOver* tool (https://genome.ucsc.edu/cgi-bin/hgLiftOver). The most representative isoform for each gene was manually selected through visualization of individual profiles.

### Splicing intermediate and lariat detection

Exons containing splicing intermediates or introns containing lariats were identified using a peak finder algorithm (NET_snrPeakFinder) (Churchman and Weissman 2011; Prudêncio et al. 2020) that detects the presence of a peak in the last nucleotide of an exon (splicing intermediate) or an intron (lariat), by comparing the accumulation of 3′ end reads mapping at that position with the mean read density of the flanking 200 nucleotides. A peak is called when the read density at the peak is superior to the mean of this surrounding region plus 3 standard deviations (Churchman and Weissman 2011). Since gene read density influences peak detection, the exons were divided into quartiles based on the *d*NET-seq read density of the corresponding gene. Only exons from the highest quartile (i.e. from genes with the highest read density) were considered in **Fig 2** for *d*NET-seq Late analysis.

### Analysis of read density and peak calling

RPKM values for the merged *d*NET-seq datasets were calculated in the following manner:

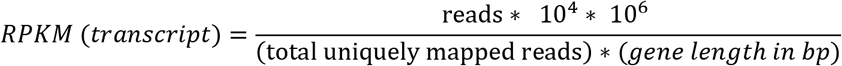

Where 10^4^ normalizes for gene length and 10^6^ normalizes for sequencing depth. Queries of gene 3’UTR overlaps (**Figure 3**) between genes were performed with bedtools intersect (version 2.27.1-1-gb87c465) (Quinlan and Hall 2010).

To detect splicing intermediate and intron lariat peaks, the algorithm from Churchman and Weissman (2011) was adapted as described in Prudêncio et al. (2020). To be able to call larger regions as peaks, a custom algorithm was developed and implemented in Python 3.7. Note that like for all of the custom Python code reported here, there is heavy reliance on *bedtools* v2.29.0 (Quinlan and Hall 2010) for operations on coordinate intervals. In addition, NumPy v1.17.2 (Harris et al. 2020) and SciPy 1.4.0 (SciPy 1.0 Contributors et al. 2020) were used. Our peak caller detects regions where the local read density is significantly higher than expected by chance given the overall read density of the transcript. Almost all of the numerical parameters used by the peak caller can be adjusted. In **Fig 4A**, results obtained with two particular parameterizations are shown. Peak Caller 1 (“large peaks”) is adapted to detecting larger peaks and provides results that are more intuitive to a human observer. Peak Caller 2 (“small peaks”) provides a finer spatial resolution, and corresponds to the settings used in all of the analyses in this study. The peak caller takes as input a BED file with the 3’ ends of reads (single-nucleotide resolution), as well as a GTF file with transcript and exon annotations and a list of transcripts to analyse. It functions by calculating a sliding average of read density within each transcript (window size 5/21 for small/large; only reads mapping to the same strand as the annotated transcript are considered). It then randomly shuffles the positions of the reads within the transcript and recalculates the sliding averages to determine the random expectation. This can be repeated several times (5 in this study) for more robustness. Windows obtained with the true read distribution are called as significant if their read density is higher than the 99^th^ percentile of the simulated windows. Note that in this study, we used a setting whereby this threshold is calculated separately for each exon (and its upstream intron, for exons other than the first), by excluding the intron-exon pair of interest and reads overlapping it during the simulation step. This is necessary so that when calling peaks within a given exon (and its upstream intron), the threshold set would not be affected by the reads within that particular exon and its upstream intron. This way, for instance, the calling of a peak in the beginning of the exon is not affected by the calling of a peak in the middle of the same exon (except through potential merging, see below). After the initial peaks are called, they are filtered to remove peaks where more than 90% of the reads come from a single nucleotide (probable PCR duplicates), that are shorter than 5 nucleotides, or that overlap with fewer reads than a specified threshold (10/5 for large/small). Finally, peaks that are within a specified distance of each-other (21/5 nucleotides for large/small) are merged together.

### Individual gene profiles

Individual dNET-seq gene profiles were generated by separating reads by strand using SAMtools (version 1.7) (Li et al. 2009). Strand-separated read data was converted to bedGraph format using bedtools genomecov with the -bg flag (version 2.27.1-1-gb87c465) (Quinlan and Hall 2010). Coverage values were normalized per nucleotide accounting for the total number of uniquely aligned reads and with the scale set to reads per 10^8^ sequences. The outcome was converted to bigwig files through the bedGraphToBigWig tool (Kent et al. 2010) and uploaded to the UCSC genome browser (James Kent et al. 2002).

### Metagene analysis

The read density metagene plots in **Fig 3F-I** and **Fig 5E,F** were created with deepTools (version 3.0.2) (Ramírez et al. 2016). Metagenes with normalized gene size (Figure 3) have bins of 10 bp while all other metagene plots in this study have single nucleotide resolution. Normalized gene and intron lengths (**Fig 3F-I**) were obtained through the scale-regions option. Exon-intron junctions without normalized lengths were obtained using the reference-option set to the 3’SS or the 5’SS. For normalization, we divided the number of reads at each nucleotide (or bin) by the total number of reads in the entire genomic region under analysis. These values were then used to calculate the mean for each nucleotide, and the results were plotted in an arbitrary units (A.U.) ranging from 0 to 1.

The peak or read density metagene plots in **Fig 3D**, **4B** and **7D**, and **Supplemental Fig S3G, S7A-G** and **S7E-J** were prepared using custom Python and R scripts (https://github.com/rosinaSav/dNETseq_code). The peak density value represents the proportion of introns/exons that overlap with a peak at that position. Only internal fully coding exons that were at least 100 nucleotides long were included. In addition, the intron just upstream of the exon had to be at least 50 nt long. For **Fig 7D**, **S7A-G** and **S7E-H**, further filtering based on read coverage was performed (see below). Note that exons shorter than 150 nucleotides contribute both to the upstream and downstream exonic proportion of the plot.

### Immediate splicing analysis

The immediate splicing analysis was performed solely on the S5P data sets. Only the 117 previously annotated pre-MBT genes (Chen et al. 2013) were analysed for early data sets. Reads were considered as spliced if they contained ‘N’s in the CIGAR string, in a position corresponding to an annotated intron. Reads that overlapped both the (unspliced) intron and the downstream exon were considered as unspliced. In both cases, only reads where the 3’ end was located at least 5 nt downstream of the 3’ ss were included, to avoid analysing misaligned reads whose 3’ end should have mapped to the end of the upstream exon instead. Spliced reads that had the 5’ end mapped to the upstream exon and the 3’ end mapped to the intron were considered indicative of recursive splicing if the first nucleotide of the downstream end (indicative of the ratchet point position) matched the second G in the AGGT canonical splicing motif. If the last nucleotide of a read matched the last nucleotide of an exon, it was considered a splicing intermediate read and not representative of nascent RNA. A splicing ratio was calculated by dividing the number of nascent RNA spliced reads by the sum of the number of spliced and unspliced nascent RNA reads, only including reads whose 3’ ends mapped to the first 100 nt of the downstream exon. Only fully coding internal exons at least 100 nt long were considered (exceptionally, in **Fig 6E** and **K** and **Supplemental Fig S6O-Q**, the 3’ most coding exon was also analysed). Finally, we performed filtering to remove exons where the read coverage was too low to allow for robust estimation of the splicing ratio. The relevant threshold was calculated for each dataset separately. We calculated the total proportion of spliced reads out of all spliced/unspliced reads for the dataset to obtain the expected splicing ratio. We then performed a binomial test to know the probability of sampling only spliced/unspliced reads by chance under the null that the true splicing ratio equalled this expectation. We set the threshold as the lowest number of reads that had to be sampled for the probability to be below 0.01. Through his procedure, the threshold was set at >=10 reads for replicate 1 and 2 of the late data set (no terminal coding exons), at >=11/10 reads for replicate 1/2 of the late data set including terminal coding exons, and at >=14/9 reads for replicate 1/2 of the early data set (terminal coding exons always included).

### Gene architecture and nucleotide composition analysis

Gene architecture and nucleotide composition parameters were calculated using custom Python and R scripts based on Ensembl annotations for dm6.18 (Cunningham et al. 2019). Splice site strength scores were calculated using MaxEntScan (Yeo and Burge 2004) with default parameters. Significance testing was performed using Kruskal-Wallis tests with a Bonferroni correction for multiple comparisons, with the correction applied separately for either replicate. For predictors where the corrected *P*-value was <0.05, Dunn’s test was performed on the pairwise comparisons using *R* package *dunn.test* (Dinno 2017). The sequence logo in **Fig 4D** was generated using the *seqLogo* package version 1.52.0 in R (Bembom 2019).

### Data access

All raw and processed sequencing data generated in this study have been submitted to the NCBI Gene Expression Omnibus (GEO; https://www.ncbi.nlm.nih.gov/geo/) under accession number GSE152585. The Python code used is available at https://github.com/rosinaSav/dNETseq_code and https://github.com/kennyrebelo. The Python code can also be found in the **Supplemental Code** file.

## Acknowledgements

We thank Takayuki Nojima and Nicholas Proudfoot (University of Oxford, UK) for critical discussion. This work was supported by funding to M.C.-F. (Fundação para a Ciência e Tecnologia, FCT/ Ministério da Ciência, Tecnologia e Ensino Superior -Fundos do Orçamento de Estado (UIDB/50005/2020), and FCT/FEDER/POR Lisboa 2020, Programa Operacional Regional de Lisboa PORTUGAL 2020, grant LISBOA-01-0145-FEDER-016394) and to R.G.M (FCT grant PTDC/BIA-BID/28441/2017). P.P. was a recipient of a FCT fellowship (SFRH/BD/109689/2015). R.S. was a recipient of an EMBO Long-Term Fellowship (EMBO ALTF 101-2019). This project has received funding from the European Union’s Horizon 2020 research and innovation programme under grant agreements No 842695 (Marie Skłodowska-Curie Actions) and No 857119 (RiboMed).

## Author Contributions

P.P. performed all molecular biology experiments. R.S. performed the majority of splicing kinetics analyses and developed the peak caller for detecting putative pause sites. K.R. performed additional bioinformatics analyses. P.P., R.S., R.G.M. and M.C.-F conceived the study and wrote the paper.

## REFERENCES

Akhtar J, Kreim N, Marini F, Mohana G, Brüne D, Binder H, Roignant J-Y. 2019. Promoter-proximal pausing mediated by the exon junction complex regulates splicing. Nat Commun 10: 521.

Alexander RD, Innocente SA, Barrass JD, Beggs JD. 2010. Splicing-dependent RNA polymerase pausing in yeast. Molecular Cell 40: 582–593.

Alpert T, Herzel L, Neugebauer KM. 2017. Perfect timing: splicing and transcription rates in living cells: Splicing and transcription rates in living cells. WIREs RNA 8: e1401.

Ameur A, Zaghlool A, Halvardson J, Wetterbom A, Gyllensten U, Cavelier L, Feuk L. 2011. Total RNA sequencing reveals nascent transcription and widespread co-transcriptional splicing in the human brain. Nature Structural and Molecular Biology 18: 1435–1440.

Artieri CG, Fraser HB. 2014. Transcript length mediates developmental timing of gene expression across drosophila. Molecular Biology and Evolution 31: 2879–2889.

Arzate-Mejía RG, Josué Cerecedo-Castillo A, Guerrero G, Furlan-Magaril M, Recillas-Targa F. 2020. In situ dissection of domain boundaries affect genome topology and gene transcription in Drosophila. Nat Commun 11: 894.

Aslanzadeh V, Huang Y, Sanguinetti G, Beggs JD. 2018. Transcription rate strongly affects splicing fidelity and cotranscriptionality in budding yeast. Genome Research 28: 203–213.

Batsché E, Yaniv M, Muchardt C. 2006. The human SWI/SNF subunit Brm is a regulator of alternative splicing. Nature Structural & Molecular Biology 13: 22–29.

Bembom O. 2019. seqLogo: Sequence logos for DNA sequence alignments. R package version 1.52.0.

Berget SM. 1995. Exon Recognition in Vertebrate Splicing. The Journal of Biological Chemistry 270: 2411–2414.

Beyer AL, Osheim YN. 1988. Splice site selection, rate of splicing, and alternative splicing on nascent transcripts. Genes & Development 2: 754–765.

Blythe SA, Wieschaus EF. 2015. Coordinating Cell Cycle Remodeling with Transcriptional Activation at the Drosophila MBT. In Current Topics in Developmental Biology, Vol. 113 of, pp. 113–148, Elsevier.

Blythe SA, Wieschaus EF. 2016. Establishment and maintenance of heritable chromatin structure during early Drosophila embryogenesis. eLife 5: e20148.

Boija A, Mahat DB, Zare A, Holmqvist P-H, Philip P, Meyers DJ, Cole PA, Lis JT, Stenberg P, Mannervik M. 2017. CBP Regulates Recruitment and Release of Promoter-Proximal RNA Polymerase II. Molecular Cell 68: 491–503.e5.

Brugiolo† M, Herzel† L, Neugebauer KM. 2013. Counting on co-transcriptional splicing. F1000Prime Rep 5: 9.

Burnette JM, Miyamoto-Sato E, Schaub MA, Conklin J, Lopez AJ. 2005. Subdivision of large introns in Drosophila by recursive splicing at nonexonic elements. Genetics 170: 661–674.

Bushnell B, Rood J, Singer E. 2017. BBMerge – Accurate paired shotgun read merging via overlap ed. P.J. Biggs. PLOS ONE 12: e0185056.

Campos-Ortega JA. 1985. Genetics of early neurogenesis in Drosophila melanogaster. Trends in Neurosciences 8: 245–250.

Carrillo Oesterreich F, Herzel L, Straube K, Hujer K, Howard J, Neugebauer KM. 2016. Splicing of Nascent RNA Coincides with Intron Exit from RNA Polymerase II. Cell 165: 372–381.

Carrillo Oesterreich F, Preibisch S, Neugebauer KM. 2010. Global analysis of nascent RNA reveals transcriptional pausing in terminal exons. Molecular Cell 40: 571–581.

Chathoth KT, Barrass JD, Webb S, Beggs JD. 2014. A Splicing-Dependent Transcriptional Checkpoint Associated with Prespliceosome Formation. Molecular Cell 53: 779–790.

Chen K, Johnston J, Shao W, Meier S, Staber C, Zeitlinger J. 2013. A global change in RNA polymerase II pausing during the Drosophila midblastula transition. eLife 2: e00861.

Churchman LS, Weissman JS. 2011. Nascent transcript sequencing visualizes transcription at nucleotide resolution. Nature 469: 368–373.

Core LJ, Waterfall JJ, Lis JT. 2008. Nascent RNA Sequencing Reveals Widespread Pausing and Divergent Initiation. Science 322: 1845–1849.

Cunningham F, Achuthan P, Akanni W, Allen J, Amode MR, Armean IM, Bennett R, Bhai J, Billis K, Boddu S, et al. 2019. Ensembl 2019. Nucleic Acids Research 47: D745–D751.

Dahlberg O, Shilkova O, Tang M, Holmqvist P-H, Mannervik M. 2015. P-TEFb, the Super Elongation Complex and Mediator Regulate a Subset of Non-paused Genes during Early Drosophila Embryo Development ed. G.P. Copenhaver. PLOS Genetics 11: e1004971.

David CJ, Boyne AR, Millhouse SR, Manley JL. 2011. The RNA polymerase II C-terminal domain promotes splicing activation through recruitment of a U2AF65-Prp19 complex. Genes and Development 25: 972–982.

de Almeida SF, Carmo-Fonseca M. 2008. The CTD role in cotranscriptional RNA processing and surveillance. FEBS Letters 582: 1971–1976.

De Conti L, Baralle M, Buratti E. 2013. Exon and intron definition in pre-mRNA splicing. Wiley interdisciplinary reviews RNA 4: 49–60.

de la Mata M, Alonso CR, Kadener S, Fededa JP, Blaustein M, Pelisch F, Cramer P, Bentley D, Kornblihtt AR. 2003. A Slow RNA Polymerase II Affects Alternative Splicing In Vivo. Molecular Cell 12: 525–532.

Dingwall C, Lomonossoff GP, Laskey RA. 1981. High sequence specificity of micrococcal nuclease. Nucleic Acids Research 9: 2659–2674.

Dinno A. 2017. dunn.test: Dunn’s Test of Multiple Comparisons Using Rank Sums. R package version 1.3.5.

Dobin A, Davis CA, Schlesinger F, Drenkow J, Zaleski C, Jha S, Batut P, Chaisson M, Gingeras TR. 2013. STAR: Ultrafast universal RNA-seq aligner. Bioinformatics 29: 15–21.

Drexler HL, Choquet K, Churchman LS. 2020. Splicing Kinetics and Coordination Revealed by Direct Nascent RNA Sequencing through Nanopores. Molecular Cell 77: 985–998.e8.

Duff MO, Olson S, Wei X, Garrett SC, Osman A, Bolisetty M, Plocik A, Celniker SE, Graveley BR. 2015. Genome-wide identification of zero nucleotide recursive splicing in Drosophila. Nature 521: 376–379.

Emili A, Shales M, McCracken S, Xie W, Tucker PW, Kobayashi R, Blencowe BJ, Ingles CJ. 2002. Splicing and transcription-associated proteins PSF and p54nrb/NonO bind to the RNA polymerase II CTD. Rna 8: 1102–1111.

Erickson JW, Cline TW. 1993. A bZIP protein, sisterless-a, collaborates with bHLH transcription factors early in Drosophila development to determine sex. Genes and Development 7: 1688–1702.

Fong N, Kim H, Zhou Y, Ji X, Qiu J, Saldi T, Diener K, Jones K, Fu XD, Bentley DL. 2014. Pre-mRNA splicing is facilitated by an optimal RNA polymerase II elongation rate. Genes and Development 28: 2663–2676.

Fox-Walsh KL, Dou Y, Lam BJ, Hung S -p., Baldi PF, Hertel KJ. 2005. The architecture of pre-mRNAs affects mechanisms of splice-site pairing. Proceedings of the National Academy of Sciences 102: 16176–16181.

Fukaya T, Lim B, Levine M. 2017. Rapid Rates of Pol II Elongation in the Drosophila Embryo. Current Biology 27: 1387–1391.

Gaffney DJ, McVicker G, Pai AA, Fondufe-Mittendorf YN, Lewellen N, Michelini K, Widom J, Gilad Y, Pritchard JK. 2012. Controls of Nucleosome Positioning in the Human Genome ed. E. Segal. PLoS Genet 8: e1003036.

Gelfman S, Cohen N, Yearim A, Ast G. 2013. DNA-methylation effect on cotranscriptional splicing is dependent on GC architecture of the exon-intron structure. Genome Research 23: 789–799.

Görnemann J, Barrandon C, Hujer K, Rutz B, Rigaut G, Kotovic KM, Faux C, Neugebauer KM, Séraphin B. 2011. Cotranscriptional spliceosome assembly and splicing are independent of the Prp40p WW domain. Rna 17: 2119–2129.

Graveley BR, Brooks AN, Carlson JW, Duff MO, Landolin JM, Yang L, Artieri CG, Baren MJ Van, Boley N, Booth BW, et al. 2011. The Developmental Transcriptome of Drosophila melanogaster. Nature 471: 473–479.

Guilgur LG, Prudêncio P, Sobral D, Liszekova D, Rosa A, Martinho RG. 2014. Requirement for highly efficient pre-mRNA splicing during Drosophila early embryonic development. eLife 3: e02181.

Harlen KM, Churchman LS. 2017. The code and beyond: transcription regulation by the RNA polymerase II carboxy-terminal domain. Nature Reviews Molecular Cell Biology 18: 263–273.

Harris CR, Millman KJ, van der Walt SJ, Gommers R, Virtanen P, Cournapeau D, Wieser E, Taylor J, Berg S, Smith NJ, et al. 2020. Array programming with NumPy. Nature 585: 357–362.

Hatton AR, Subramaniam V, Lopez AJ. 1998. Generation of alternative Ultrabithorax isoforms and stepwise removal of a large intron by resplicing at exon-exon junctions. Molecular Cell 2: 787–796.

Herzel L, Straube K, Neugebauer KM. 2018. Long-read sequencing of nascent RNA reveals coupling among RNA processing events. 1008–1019.

Hörz W, Altenburger W. 1981. Sequence specific cleavage of DNA by micrococcal nuclease. Nucleic Acids Research 9: 2643–2658.

Hoskins RA, Landolin JM, Brown JB, Sandler JE, Takahashi H, Lassmann T, Booth BW, Zhang D, Wan KH, Yang L, et al. 2011. Genome-wide analysis of promoter architecture in Drosophila melanogaster. Genome Research 21: 182–192.

Hsin JP, Manley JL. 2012. The RNA polymerase II CTD coordinates transcription and RNA processing. Genes and Development 26: 2119–2137.

James Kent W, Sugnet CW, Furey TS, Roskin KM, Pringle TH, Zahler AM, Haussler D. 2002. The human genome browser at UCSC. Genome Research 12: 996–1006.

Ji J-Y, Squirrell JM, Schubiger G. 2004. Both Cyclin B levels and DNA-replication checkpoint control the early embryonic mitoses in Drosophila Jun-Yuan. Development 131: 401–411.

Jonkers I, Kwak H, Lis JT. 2014. Genome-wide dynamics of Pol II elongation and its interplay with promoter proximal pausing, chromatin, and exons. eLife 3: e02407.

Joseph B, Kondo S, Lai EC. 2018. Short cryptic exons mediate recursive splicing in Drosophila. Nature Structural and Molecular Biology 25: 365–371.

Kent WJ, Zweig AS, Barber G, Hinrichs AS, Karolchik D. 2010. BigWig and BigBed: Enabling browsing of large distributed datasets. Bioinformatics 26: 2204–2207.

Keren H, Lev-Maor G, Ast G. 2010. Alternative splicing and evolution: Diversification, exon definition and function. Nature Reviews Genetics 11: 345–355.

Khodor YL, Menet JS, Tolan M, Rosbash M. 2012. Cotranscriptional splicing efficiency differs dramatically between Drosophila and mouse. RNA 18: 2174–2186.

Khodor YL, Rodriguez J, Abruzzi KC, Tang CHA, Marr MT, Rosbash M. 2011. Nascent-seq indicates widespread cotranscriptional pre-mRNA splicing in Drosophila. Genes and Development 25: 2502–2512.

Kwak H, Fuda NJ, Core LJ, Lis JT. 2013. Precise maps of RNA polymerase reveal how promoters direct initiation and pausing. Science 339: 950–953.

Kwasnieski JC, Orr-Weaver TL, Bartel DP. 2019. Early genome activation in Drosophila is extensive with an initial tendency for aborted transcripts and retained introns. Genome Research 29: 1188–1197.

Larson MH, Mooney RA, Peters JM, Windgassen T, Nayak D, Gross CA, Block SM, Greenleaf WJ, Landick R, Weissman JS. 2014. A pause sequence enriched at translation start sites drives transcription dynamics in vivo. Science 344: 1042–1047.

Laver JD, Marsolais AJ, Smibert CA, Lipshitz HD. 2015. Regulation and Function of Maternal Gene Products During the Maternal-to-Zygotic Transition in Drosophila. 1st ed. Elsevier Inc.

Lee JEA, Mitchell NC, Zaytseva O, Chahal A, Mendis P, Cartier-Michaud A, Parsons LM, Poortinga G, Levens DL, Hannan RD, et al. 2015. Defective Hfp-dependent transcriptional repression of dMYC is fundamental to tissue overgrowth in Drosophila XPB models. Nat Commun 6: 7404.

Li H, Handsaker B, Wysoker A, Fennell T, Ruan J, Homer N, Marth G, Abecasis G, Durbin R. 2009. The Sequence Alignment/Map format and SAMtools. Bioinformatics 25: 2078–2079.

Lott SE, Villalta JE, Schroth GP, Luo S, Tonkin LA, Eisen MB. 2011. Noncanonical Compensation of Zygotic X Transcription in Early Drosophila melanogaster Development Revealed through Single-Embryo RNA-Seq ed. R.S. Hawley. PLoS Biology 9: e1000590.

Lund E, Dahlberg JE. 1992. Cyclic 2 ‘, 3 “-Phosphates and Nontemplated Nucleotides at the 3” End of Spliceosomal U6 Small Nuclear RNAs. Science 255: 327–330.

Martin M. 2011. Cutadapt removes adapter sequences from high-throughput sequencing reads. EMBnet.journal 17: 10–12.

Martin RM, Rino J, Carvalho C, Kirchhausen T, Carmo-Fonseca M. 2013. Live-Cell Visualization of Pre-mRNA Splicing with Single-Molecule Sensitivity. Cell Reports 4: 1144–1155.

Martinho RG, Guilgur LG, Prudêncio P. 2015. How gene expression in fast-proliferating cells keeps pace. BioEssays 37: 514–524.

Maslon MM, Braunschweig U, Aitken S, Mann AR, Kilanowski F, Hunter CJ, Blencowe BJ, Kornblihtt AR, Adams IR, Cáceres JF. 2019. A slow transcription rate causes embryonic lethality and perturbs kinetic coupling of neuronal genes. EMBO J 38: e101244.

Mayer A, Di Iulio J, Maleri S, Eser U, Vierstra J, Reynolds A, Sandstrom R, Stamatoyannopoulos JA, Churchman LS. 2015. Native elongating transcript sequencing reveals human transcriptional activity at nucleotide resolution. Cell 161: 541–554.

Morris DP, Greenleaf AL. 2000. The splicing factor, Prp40, binds the phosphorylated carboxyl-terminal domain of RNA Polymerase II. Journal of Biological Chemistry 275: 39935–39943.

Nojima T, Gomes T, Carmo-Fonseca M, Proudfoot NJ. 2016. Mammalian NET-seq analysis defines nascent RNA profiles and associated RNA processing genome-wide. Nature Protocols 11: 413–428.

Nojima T, Gomes T, Grosso ARF, Kimura H, Dye MJ, Dhir S, Carmo-Fonseca M, Proudfoot NJ. 2015. Mammalian NET-seq reveals genome-wide nascent transcription coupled to RNA processing. Cell 161: 526–540.

Nojima T, Rebelo K, Gomes T, Grosso AR, Proudfoot NJ, Carmo-Fonseca M. 2018. RNA Polymerase II Phosphorylated on CTD Serine 5 Interacts with the Spliceosome during Co-transcriptional Splicing. Molecular Cell 72: 369–379.e4.

Pai AA, Henriques T, McCue K, Burkholder A, Adelman K, Burge CB. 2017. The kinetics of pre-mRNA splicing in the Drosophila genome and the influence of gene architecture. eLife 6: e32537.

Pai AA, Paggi JM, Yan P, Adelman K, Burge CB. 2018. Numerous recursive sites contribute to accuracy of splicing in long introns in flies ed. H.D. Madhani. PLOS Genetics 14: e1007588.

Pertea M, Pertea GM, Antonescu CM, Chang TC, Mendell JT, Salzberg SL. 2015. StringTie enables improved reconstruction of a transcriptome from RNA-seq reads. Nature Biotechnology 33: 290–295.

Pritchard DK, Schubiger G. 1996. Activation of transcription in Drosophila embryos is a gradual process mediated by the nucleocytoplasmic ratio. Genes and Development 10: 1131–1142.

Proudfoot NJ. 2016. Transcriptional termination in mammals: Stopping the RNA polymerase II juggernaut. Science 352: aad9926.

Prudêncio P, Rebelo K, Grosso AR, Martinho RG, Carmo-fonseca M. 2020. Analysis of Mammalian Native Elongating Transcript sequencing (mNET-seq)high-throughput data. Methods 178: 89–95.

Quinlan AR, Hall IM. 2010. BEDTools: A flexible suite of utilities for comparing genomic features. Bioinformatics 26: 841–842.

Ramírez F, Ryan DP, Grüning B, Bhardwaj V, Kilpert F, Richter AS, Heyne S, Dündar F, Manke T. 2016. deepTools2: a next generation web server for deep-sequencing data analysis. Nucleic acids research 44: W160–W165.

Reimer KA, Mimoso CA, Adelman K, Neugebauer KM. 2021. Co-transcriptional splicing regulates 3′ end cleavage during mammalian erythropoiesis. Molecular Cell 81: 998–1012.e7.

Robberson BL, Cote GJ, Berget SM. 1990. Exon Definition May Facilitate Splice Site Selection in RNAs with Multiple Exons. Molecular and Cellular Biology 10: 84–94.

Rothe M, Pehl M, Taubert H, Jäckle H. 1992. Loss of gene function through rapid mitotic cycles in the Drosophila embryp. Nature 359: 156–159.

Sandler JE, Irizarry J, Stepanik V, Dunipace L, Amrhein H, Stathopoulos A. 2018. A Developmental Program Truncates Long Transcripts to Temporally Regulate Cell Signaling. Developmental Cell 47: 773–784.e6.

Saunders A, Core LJ, Sutcliffe C, Lis JT, Ashe HL. 2013. Extensive polymerase pausing during Drosophila axis patterning enables high-level and pliable transcription. Genes and Development 27: 1146–1158.

Schlackow M, Nojima T, Gomes T, Dhir A, Carmo-Fonseca M, Proudfoot NJ. 2017. Distinctive Patterns of Transcription and RNA Processing for Human lincRNAs. Molecular Cell 65: 25–38.

Schulz KN, Bondra ER, Moshe A, Villalta JE, Lieb JD, Kaplan T, McKay DJ, Harrison MM. 2015. Zelda is differentially required for chromatin accessibility, transcription factor binding, and gene expression in the early Drosophila embryo. Genome Res 25: 1715–1726.

SciPy 1.0 Contributors, Virtanen P, Gommers R, Oliphant TE, Haberland M, Reddy T, Cournapeau D, Burovski E, Peterson P, Weckesser W, et al. 2020. SciPy 1.0: fundamental algorithms for scientific computing in Python. Nat Methods 17: 261–272.

Sheridan RM, Fong N, D’Alessandro A, Bentley DL. 2019. Widespread Backtracking by RNA Pol II Is a Major Effector of Gene Activation, 5′ Pause Release, Termination, and Transcription Elongation Rate. Molecular Cell 73: 107–118.e4.

Shermoen AW, O’Farrell PH. 1991. Progression of the Cell Cycle through Mitosis Leads to Abortion of Nascent Transcripts. Cell 67: 303–310.

Sousa-Luís R, Dujardin G, Zukher I, Kimura H, Weldon C, Carmo-Fonseca M, Proudfoot NJ, Nojima T. 2021. POINT technology illuminates the processing of polymerase-associated intact nascent transcripts. Molecular Cell 81: 1935–1950.e6.

Sterner DA, Carlo T, Berget SM. 1996. Architectural limits on split genes. Proc Natl Acad Sci USA 93: 15081–15085.

Veloso A, Kirkconnell KS, Magnuson B, Biewen B, Paulsen MT, Wilson TE, Ljungman M. 2014. Rate of elongation by RNA polymerase II is associated with specific gene features and epigenetic modifications. Genome Research 24: 896–905.

Vizcaya-Molina E, Klein CC, Serras F, Mishra RK, Guigó R, Corominas M. 2018. Damage-responsive elements in *Drosophila* regeneration. Genome Res 28: 1852–1866.

Wan Y, Anastasakis DG, Rodriguez J, Palangat M, Gudla P, Zaki G, Tandon M, Pegoraro G, Chow CC, Hafner M, et al. 2021. Dynamic imaging of nascent RNA reveals general principles of transcription dynamics and stochastic splice site selection. Cell 184: 2878–2895.e20.

Windhager L, Bonfert T, Burger K, Ruzsics Z, Krebs S, Kaufmann S, Malterer G, L’Hernault A, Schilhabel M, Schreiber S, et al. 2012. Ultrashort and progressive 4sU-tagging reveals key characteristics of RNA processing at nucleotide resolution. Genome Research 22: 2031–2042.

Yan D, Neumüller RA, Buckner M, Ayers K, Li H, Hu Y, Yang-Zhou D, Pan L, Wang X, Kelley C, et al. 2014. A Regulatory Network of Drosophila Germline Stem Cell Self-Renewal. Developmental Cell 28: 459–473.

Yeo G, Burge CB. 2004. Maximum entropy modeling of short sequence motifs with applications to RNA splicing signals. Journal of Computational Biology 11: 377–394.

Yuan K, Seller CA, Shermoen AW, Farrell PHO. 2016. Timing the Drosophila Mid-Blastula Transition: A Cell Cycle-Centered View. Trends in Genetics 32: 496–507.

Zhu L, Zhang Y, Zhang W, Yang S, Chen J-Q, Tian D. 2009. Patterns of exon-intron architecture variation of genes in eukaryotic genomes. BMC Genomics 10: 47.

