## Supplemental Fig S1 for "Transcription and splicing dynamics during early *Drosophila* development"

A

| Dataset | Raw reads | Uniquely Mapped after Merge |
|---|---|---|
| Early_2-3h_S5P_rep1 | 62,891,575 | 12,959,070 |
| Early_2-3h_S5P_rep2 | 56,917,697 | 3,610,607 |
| Late_4-6h_S5P_rep1 | 65,865,258 | 21,564,673 |
| Late_4-6h_S5P_rep2 | 75,909,131 | 27,375,998 |
| Late_4-6h_S5P_rep3 | 71,923,876 | 11,987,034 |
| Late_4-6h_S2P_rep1 | 77,838,463 | 38,595,780 |
| Late_4-6h_S2P_rep2 | 64,862,996 | 23,137,529 |
| Late_4-6h_S2P_rep3 | 76,719,886 | 17,912,504 |

B

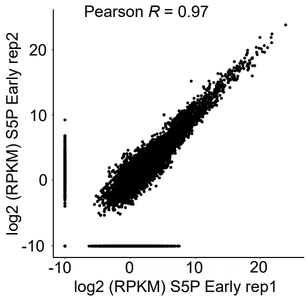


C

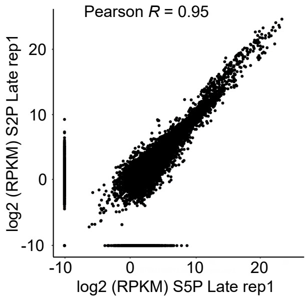


D

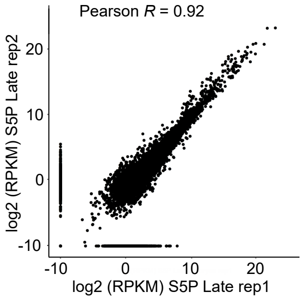


E

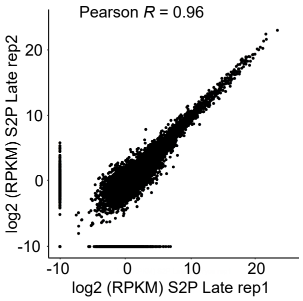


F

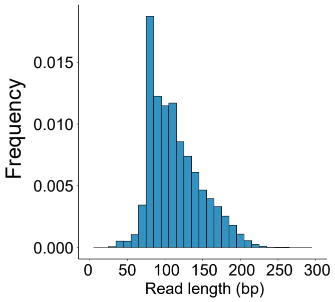


G

| Dataset | Number of nascent reads | Nascent reads / Uniquely mapped reads (%) |
|---|---|---|
| Early_2-3h_S5P_rep1 | 7,984,411 | 61.6 |
| Early_2-3h_S5P_rep2 | 2,430,876 | 67.3 |
| Late_4-6h_S5P_rep1 | 13,413,045 | 62.2 |
| Late_4-6h_S5P_rep2 | 13,777,880 | 50.3 |
| Late_4-6h_S5P_rep3 | 8,505,056 | 71.0 |
| Late_4-6h_S2P_rep1 | 23,153,659 | 60.0 |
| Late_4-6h_S2P_rep2 | 12,926,707 | 55.9 |
| Late_4-6h_S2P_rep3 | 10,356,743 | 57.8 |

H

| Dataset | min | mean | max | 25th percentile | 50th percentile | 75th percentile |
|---|---|---|---|---|---|---|
| Early_2-3h_S5P_rep1 | 35 | 99.6 | 293 | 74 | 81 | 119 |
| Early_2-3h_S5P_rep2 | 35 | 94.3 | 293 | 72 | 81 | 110 |
| Late_4-6h_S5P_rep1 | 35 | 114.2 | 293 | 86 | 108 | 135 |
| Late_4-6h_S5P_rep2 | 35 | 102.0 | 294 | 80 | 87 | 119 |
| Late_4-6h_S5P_rep3 | 35 | 88.4 | 294 | 76 | 81 | 89 |
| Late_4-6h_S2P_rep1 | 35 | 107.0 | 294 | 81 | 96 | 128 |
| Late_4-6h_S2P_rep2 | 35 | 111.1 | 294 | 81 | 102 | 132 |
| Late_4-6h_S2P_rep3 | 35 | 109.0 | 293 | 83 | 105 | 130 |

**Supplemental Fig. S1.** (A) For each *d*NET-seq library prepared, the number of total reads and of uniquely aligned reads is indicated. (B) Density of uniquely aligned reads per gene (RPKM in log2 scale) for two *d*NET-seq/S5P biological replicates from early embryos (Pearson's correlation, R = 0.97). (C) Density of uniquely aligned reads per gene (RPKM in log2 scale) from late embryos analysed by *d*NET-seq/S5P and *d*NET-seq/S2P (Pearson's correlation, R = 0.95). (D, E) Density of uniquely aligned reads per gene (RPKM in log2 scale) for two *d*NET-seq/S5P and *d*NET-seq/S2P biological replicates from late embryos (Pearson's correlation, R = 0.92; R = 0.96). (F) Histogram of the read lengths in *d*NET-seq/S5P data from late embryos. (G) For each *d*NET-seq library generated, the number of uniquely aligned reads corresponding to nascente transcripts is indicated. (H) Length of sequenced nascent RNA (in nucleotides).
