## Supplemental Fig S2 for "Transcription and splicing dynamics during early *Drosophila* development"

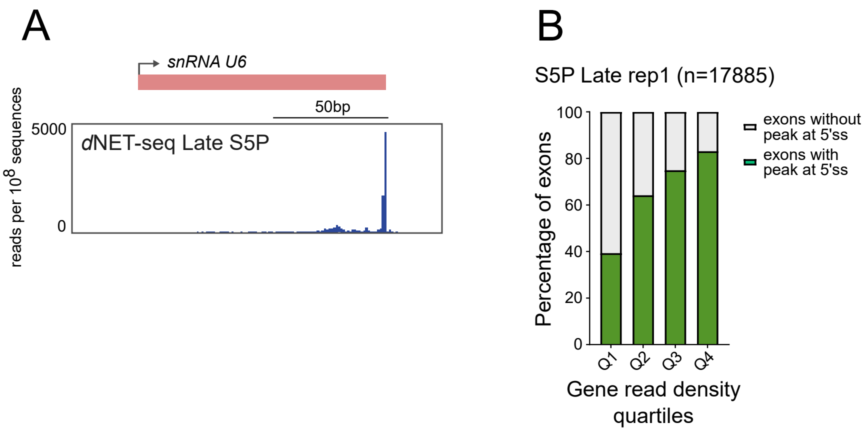


**Supplemental Fig. S2.** (**A**) *d*NET-seq/S5P profile over the U6 snRNA gene in the late dataset (replicate 1). (**B**) Frequency of peaks corresponding to splicing intermediates (green) detected by *d*NET-seq/S5Pon exons of genes expressed in late embryos. Genes were grouped into quartiles (Q) based on their *d*NET-seqread density.
