## Supplemental Fig S3 for "Transcription and splicing dynamics during early *Drosophila* development"

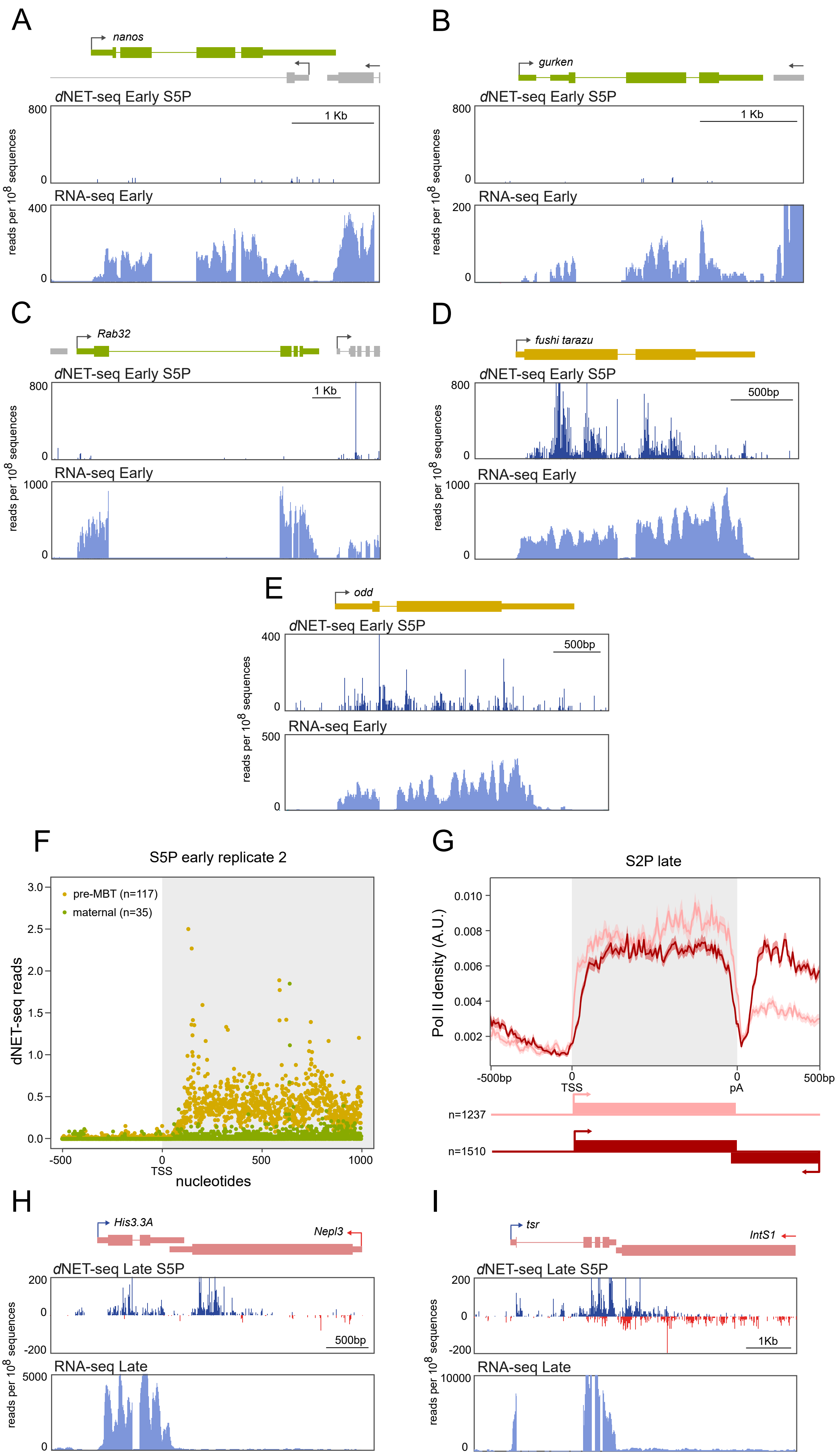

A
nanos
dNET-seq Early S5P
800
0
1 Kb
RNA-seq Early
400
0
reads per 10^8 sequences
B
gurken
dNET-seq Early S5P
800
0
1 Kb
RNA-seq Early
200
0
reads per 10^8 sequences
C
Rab32
dNET-seq Early S5P
800
0
1 Kb
RNA-seq Early
1000
0
reads per 10^8 sequences
D
fushi tarazu
dNET-seq Early S5P
800
0
500bp
RNA-seq Early
1000
0
reads per 10^8 sequences
E
odd
dNET-seq Early S5P
400
0
500bp
RNA-seq Early
500
0
reads per 10^8 sequences
F
S5P early replicate 2
pre-MBT (n=117)
maternal (n=35)
dNET-seq reads
3.0
2.5
2.0
1.5
1.0
0.5
0.0
-500
0
TSS
500
1000
nucleotides
G
S2P late
Pol II density (A.U.)
0.010
0.008
0.006
0.004
0.002
-500bp
0
TSS
0
pA
500bp
n=1237
n=1510
H
His3.3A
Nepl3
dNET-seq Late S5P
200
0
-200
500bp
RNA-seq Late
5000
0
reads per 10^8 sequences
I
tsr
IntS1
dNET-seq Late S5P
200
0
-200
1Kb
RNA-seq Late
10000
0
reads per 10^8 sequences

**Supplemental Fig. S3.** (A-C) *d*NET-seq/S5P and RNA-seq profiles over maternal genes in early embryos: *nanos* (A), *gurken* (B), and *Rab32* (C). (D-E) *d*NET-seq/S5P and RNA-seq profiles over pre-MBT genes in early embryos: (D) *fushi tarazu* and *odd skipped* (*odd*) (E). (F) Meta-analysis of *d*NET-seq/S5P mean read density around the transcription start site (TSS) in maternal and pre-MBT genes (replicate 2). (G) Normalized metagene analysis in arbitrary units (A.U.). The *d*NET-seq/S2P signal over transcriptionally active genes in late embryos is depicted along the normalized gene length (grey background), as well as 500 bp upstream of the transcription start site (TSS) and 500 bp downstream of the polyadenylation (pA) site. The signal over genes that have the 3' UTR overlapped by an antisense gene is depicted in dark red, while the signal over genes with no other genes within 500 bp is depicted in light red. (H-I) *d*NET-seq/S5P and RNA-seq profiles over MBT genes in late embryos: *His3.3A* (H) and *twinstar* (*tsr*) (I). Reads that aligned to the positive strand are in blue, and reads that aligned to the negative strand are in red. The direction of transcription is indicated by an arrow.
