## Supplemental Fig S4 for "Transcription and splicing dynamics during early *Drosophila* development"

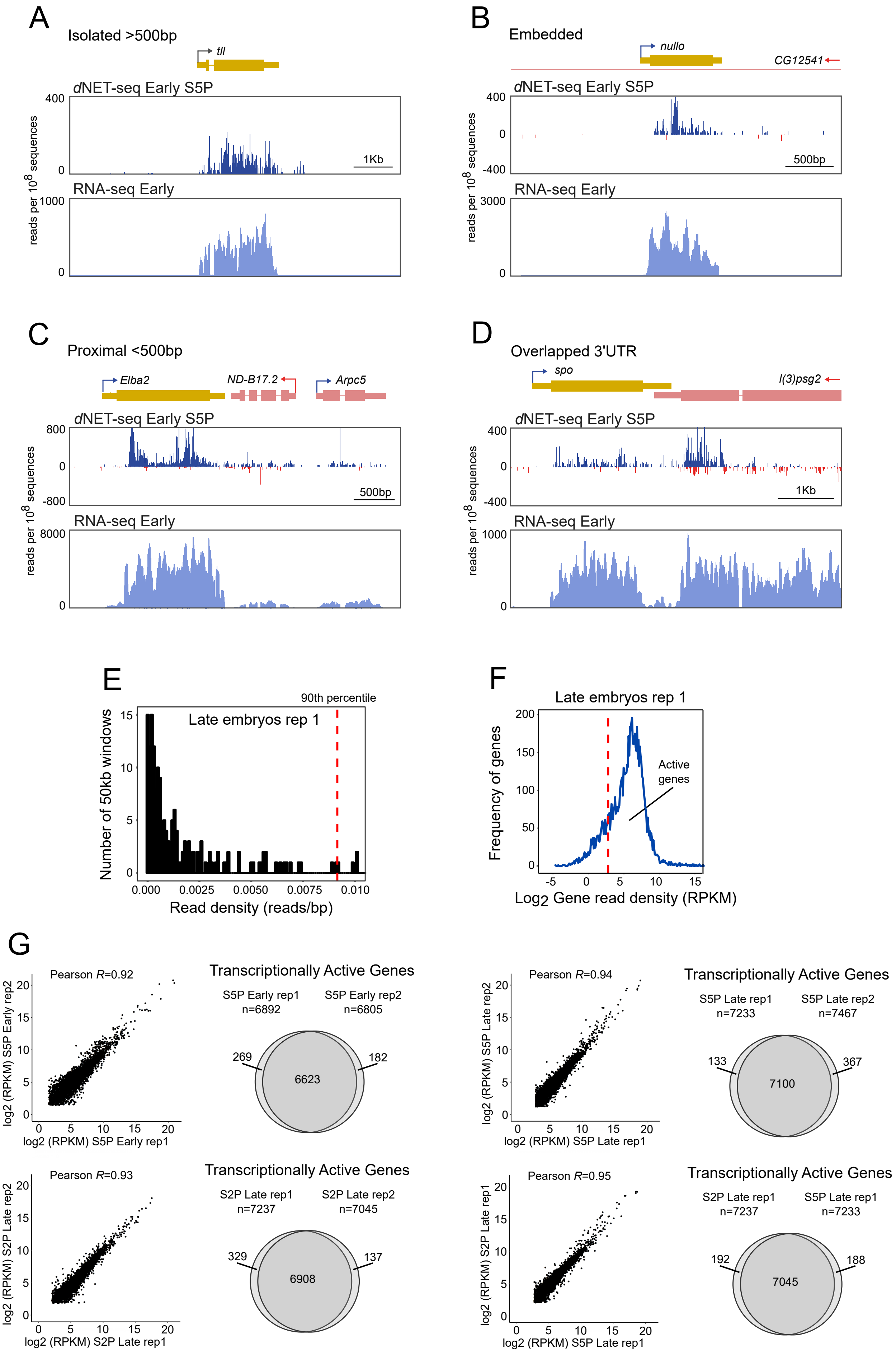

A
Isolated >500bp
tll
dNET-seq Early S5P
400
0
1Kb
RNA-seq Early
1000
0
reads per 10^8 sequences
B
Embedded
nullo
CG12541
dNET-seq Early S5P
400
0
-400
500bp
RNA-seq Early
3000
0
reads per 10^8 sequences
C
Proximal <500bp
Elba2
ND-B17.2
Arpc5
dNET-seq Early S5P
800
0
-800
500bp
RNA-seq Early
8000
0
reads per 10^8 sequences
D
Overlapped 3'UTR
spo
l(3)psg2
dNET-seq Early S5P
400
0
-400
1Kb
RNA-seq Early
1000
0
reads per 10^8 sequences
E
90th percentile
Late embryos rep 1
Number of 50kb windows
15
10
5
0
0.000
0.0025
0.0050
0.0075
0.010
Read density (reads/bp)
F
Late embryos rep 1
200
150
100
50
0
Frequency of genes
Active genes
-5
0
5
10
15
Log2 Gene read density (RPKM)
G
Pearson R=0.92
log2 (RPKM) S5P Early rep2
log2 (RPKM) S5P Early rep1
Transcriptionally Active Genes
S5P Early rep1 n=6892
S5P Early rep2 n=6805
269
6623
182
Pearson R=0.94
log2 (RPKM) S5P Late rep2
log2 (RPKM) S5P Late rep1
Transcriptionally Active Genes
S5P Late rep1 n=7233
S5P Late rep2 n=7467
133
7100
367
Pearson R=0.93
log2 (RPKM) S2P Late rep2
log2 (RPKM) S2P Late rep1
Transcriptionally Active Genes
S2P Late rep1 n=7237
S2P Late rep2 n=7045
329
6908
137
Pearson R=0.95
log2 (RPKM) S2P Late rep1
log2 (RPKM) S5P Late rep1
Transcriptionally Active Genes
S2P Late rep1 n=7237
S5P Late rep1 n=7233
192
7045
188

**Supplemental Fig. S4.** (A-D) *d*NET-seq/S5P and RNA-seq profiles over the indicated pre-MBT genes. Reads that aligned to the positive strand are in blue, and reads that aligned to the negative strand are in red. The direction of transcription is indicated by an arrow. (E-G) Identification of transcriptionally active genes based on *d*NET-seq signal along the gene body in late embryos. (E) Read density distribution in intergenic regions (gene deserts). The red dashed line represents the 90th percentile of read density in all intergenic regions analysed. (F) Read density distribution per gene (RPKM) represented in Log2 scale. The 90th percentile of read density over gene deserts is set as threshold (red dashed line). (G) Comparison of read density per active gene (RPKM in log2 scale) between the indicated datasets. Total number of active genes identified in each dataset is indicated (n). Venn diagrams show common active genes between datasets.
