## Supplemental Fig S5 for "Transcription and splicing dynamics during early *Drosophila* development"

A

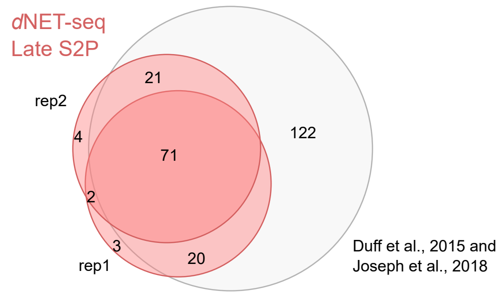


**Supplemental Fig . S5**. **(A)** Venn diagram comparing RPs identified in two *d*NET-seq/S2P biological replicates and in previously reported studies (Joseph et al. 2018; Duff et al. 2015).
