## Supplemental Fig S6 for "Transcription and splicing dynamics during early *Drosophila* development"

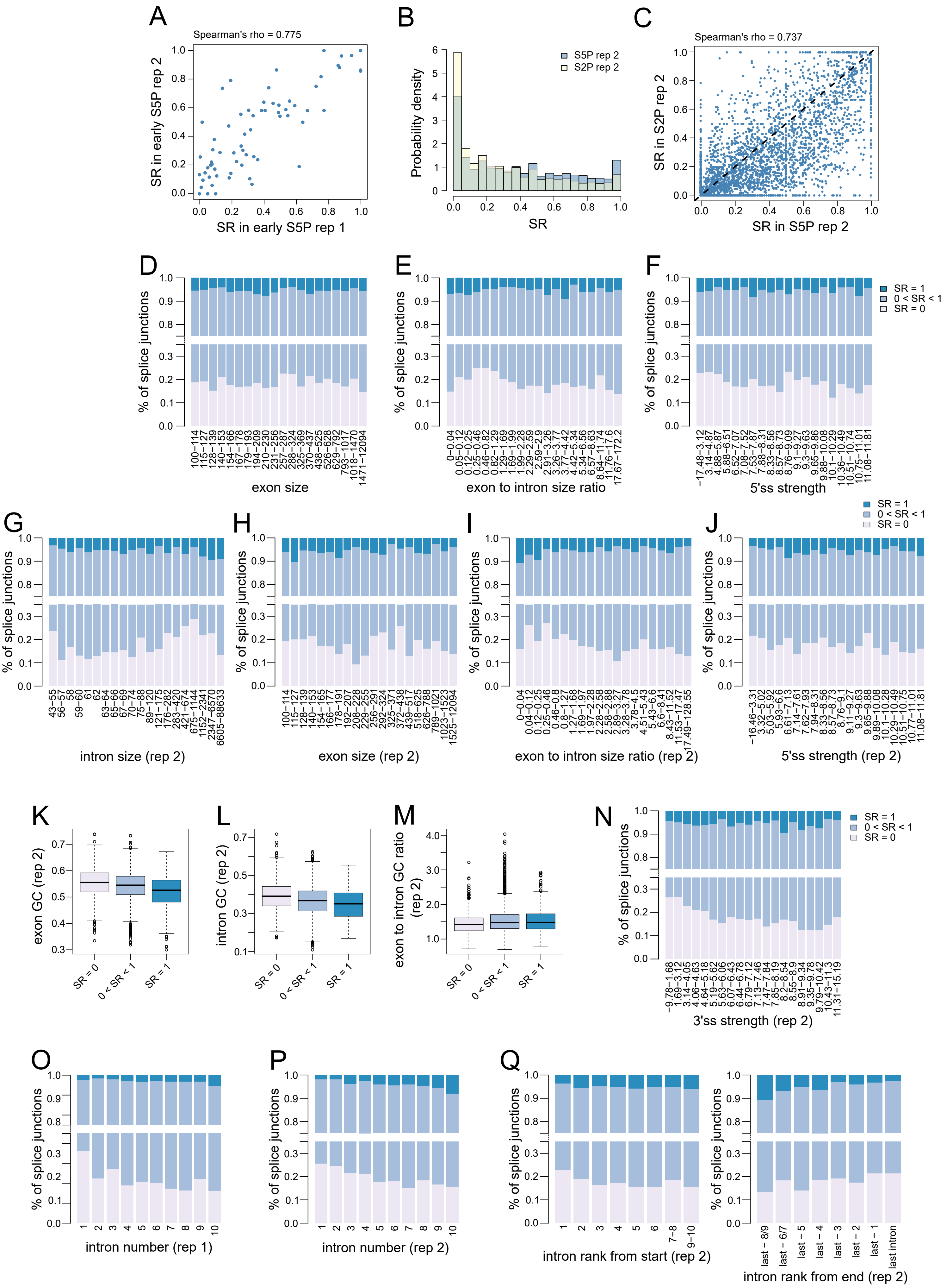

A
Spearman's rho = 0.775
SR in early S5P rep 2
SR in early S5P rep 1
B
S5P rep 2
S2P rep 2
Probability density
SR
C
Spearman's rho = 0.737
SR in S2P rep 2
SR in S5P rep 2
D
E
F
% of splice junctions
SR = 1
0 < SR < 1
SR = 0
exon size
exon to intron size ratio
5'ss strength
G
H
I
J
intron size (rep 2)
exon size (rep 2)
exon to intron size ratio (rep 2)
5'ss strength (rep 2)
K
L
M
N
exon GC (rep 2)
intron GC (rep 2)
exon to intron GC ratio (rep 2)
SR = 0
0 < SR < 1
SR = 1
3'ss strength (rep 2)
O
P
Q
intron number (rep 1)
intron number (rep 2)
intron rank from start (rep 2)
intron rank from end (rep 2)

**Supplemental Fig. S6.** In all panels, unless otherwise specified, only introns from transcriptionally active genes where the downstream exon is a fully coding internal exon at least 100 nt long were included. In addition, enough spliced/unspliced reads had to end within the first 100 exonic nucleotides that obtaining a splicing ratio (SR) of 0 or 1 by chance alone was highly unlikely (see the Methods section for details). For genes expressed in late embryos, this threshold was 10 reads for both replicates. For pre-MBT genes, it was 14 for replicate 1 and 9 for replicate 2. For S2P, a threshold of 10 was used for better comparability with S5P data. (A) SR values estimated in two biological replicates of *d*NET-seq/S5P datasets for pre-MBT genes in early embryos (Spearman correlation, ρ = ~0.775, P ~ 1.348 * 10-14). (B) Distribution of SR values for *d*NET-seq/S5P (N = 4510) and *d*NET-seq/S2P (N = 4373). (C) SR values estimated in replicate 2 of *d*NET-seq/S5P and *d*NET-seq/S2P from late embryos (Spearman correlation, ρ = ~0.737, P < 2.2 * 10-16; N = 3295). (D-Q) SR values for *d*NET seq/S5P data (replicate 1 and 2) plotted out against several gene architecture parameters. For sample sizes and statistical tests, see Supplementary Table 1. In (D-J, N-Q), the bin ranges have been set so that intron numbers would be as equal possible between bins.
