## Supplemental Fig S7 for "Transcription and splicing dynamics during early *Drosophila* development"

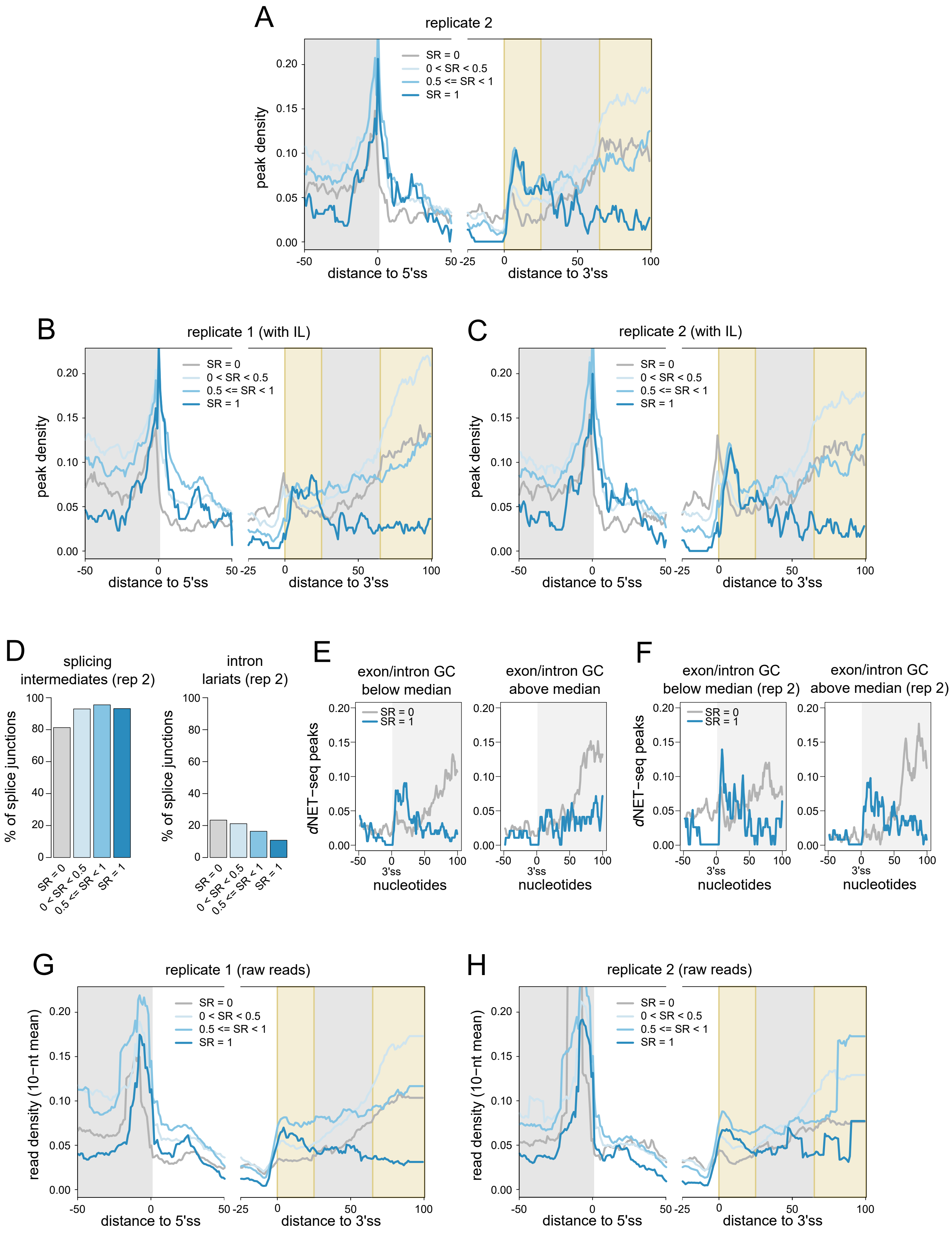
A
replicate 2
SR = 0
0 < SR < 0.5
0.5 <= SR < 1
SR = 1
peak density
distance to 5'ss
distance to 3'ss
B
replicate 1 (with IL)
C
replicate 2 (with IL)
D
splicing intermediates (rep 2)
intron lariats (rep 2)
% of splice junctions
E
exon/intron GC below median
exon/intron GC above median
dNET−seq peaks
3'ss
nucleotides
F
exon/intron GC below median (rep 2)
exon/intron GC above median (rep 2)
G
replicate 1 (raw reads)
read density (10−nt mean)
H
replicate 2 (raw reads)

**Supplemental Fig. S7.** (A-C) Metagene analyses of peak density estimated from *d*NET-seq/S5P datasets from late embryos for different SR ranges. (A) shows replicate 2, with introns with putative intron lariat reads removed (N = 3623). (B) and (C) show replicates 1 and 2, respectively, with introns with putative intron lariat reads included (N = 5614 for replicate 1 and N = 4503 for replicate 2). Peak density has been calculated as the proportion of introns that overlap with a peak at any given position. The last 50 nucleotides of exons, the first 50 nucleotides of introns, the last 25 nucleotides of introns, and the first 100 nucleotides of exons are shown. Only internal and fully coding exons from transcriptionally active genes that are at least 100 nucleotides long are shown. In addition, at least 10 spliced/unspliced reads had to end within the first 100 nucleotides of the exon. (D) The proportion of introns with at least one read whose 3' end maps to the final position of the upstream exon (putative splicing intermediates) or to the final position of the intron (putative intron lariats) in *d*NET-seq/S5P late replicate 2. (E-F) *d*NET-seq/S5P peak density profiles for late embryos, replicate 1 (E) and replicate 2 (F). The final 50 nucleotides of the intron and the first 100 nucleotides of the downstream exon are shown. For each data point, the GC content over the first 100 exonic nucleotides was divided by the GC content over the last 50 intronic nucleotides, generating the exonic to intronic GC ratio. Introns were split into two classes along the median of this metric. Only introns with an SR of 0 or 1 were plotted. (G-H) Similar to 7D and S7A, respectively, but showing read densities (averaged over a 10-nucleotide sliding window) rather than peak densities. (I-J) Metagene analysis of peak density estimated from *d*NET-seq/S2P datasets from late embryos, in replicate 1(I) and replicate 2 (J). The last 50 nucleotides of exons, the first 50 nucleotides of introns, the last 25 nucleotides of introns, and the first 100 nucleotides of exons are shown. Only internal and fully coding exons from transcriptionally active genes that are at least 100 nucleotides long are shown (N = 13229 for replicate 1, N = 11831 for replicate 2). Exons shorter than 150 nucleotides contribute to both the exon end and start. Only introns that were at least 50 nt long were considered.
