## Supplemental Methods for "Transcription and splicing dynamics during early *Drosophila* development"

### SUPPLEMENTARY METHODS

#### *d*NET-seq sequence biases

Previous work has shown that MNase cleaves DNA more frequently at AT than at GC base pairs (Dingwall *et al.*, 1981). More specifically, Hörz and Altenburger (1981) found a preference for sites where a few alternating AT-TA base pairs were surrounded by a more GC-rich region, rather than for sites within longer AT-rich stretches. More recent genome-wide work in humans has come to similar conclusions (Gaffney *et al.*, 2012). *d*NET-seq relies on MNase cleavage of both DNA and RNA to solubilize chromatin. Any nucleotide biases in cleavage may be problematic, as they could mean that the solubilization is more efficient in regions where the nucleotide composition is more in line with such cleavage biases. This could lead to an artefactually increased read density in such regions.

To verify whether there was any evidence for RNA cleavage biases, we determined the nucleotide composition around the locations of the 5’ ends of NET-seq reads (**Figure 4D** in main text). A clear preference for cleavage just 5’ of an adenine was observed, with additional weaker biases at surrounding sites. Our results are therefore broadly in line with those obtained previously for DNA.

To what extent could these biases affect our results? We generated a simulated version of a *d*NET-seq dataset (S5P late, replicate 1) where the read positions were randomized in such a way as to preserve the nucleotide biases at the 5’ ends of reads. Only reads that overlapped transcriptionally active genes were considered. More specifically, for each read, we defined the “starting hexamer” as the hexamer centred on the 5’ end of the read (the three nucleotides just 5’ of the read and the first three nucleotides of the read). We then randomly picked another occurrence of the same hexamers from within transcriptionally active genes and defined a “simulated read” at that position, preserving the length of the initial read. We then called peaks on the simulated reads similarly to true reads.

The mean proportion of nucleotides within simulated peaks was similar for both exons and introns (~0.017). However, this belies a difference in medians, with a higher peak density in exons than in introns (exon median: ~0.008; intron median: 0; $P < 2.2 * 10^{-16}$; two-tailed Mann-Whitney *U*-test; **Supplementary Methods Figure 1**). To know if the 5’ end nucleotide bias effect could explain the bias towards exons observed in real data, we calculated a normalized peak density for each exon and intron, as $\frac{true\ peak\ density - simulated\ peak\ density}{simulated\ peak\ density}$. Exons and introns with a simulated peak density of 0 were discarded. Normalized peak densities were still higher for exons than for introns ($P < 2.2 * 10^{-16}$; two-tailed Mann-Whitney *U*-test). Hence, 5’ end nucleotide biases may exaggerate but do not explain the bias towards exons in real data.

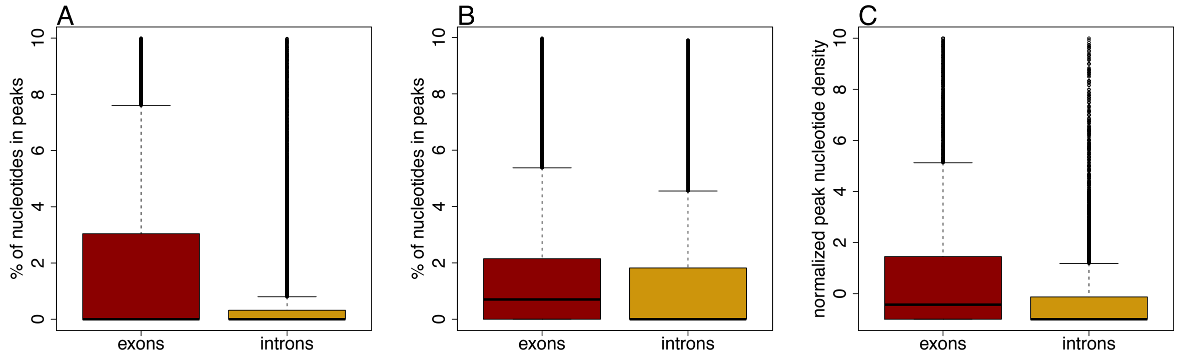


*Supplementary Methods Figure 1: distribution of peak densities for exons and introns. A: true peaks. B: simulated peaks. C: normalized peaks, obtained as* $\frac{\text{true peak density} - \text{simulated peak density}}{\text{simulated peak density}}$*. For all three panels, the y axis has been limited to values below 10 for visualization reasons.*

It may seem surprising that simulated read density would be higher in exons, given that *Drosophila* exons tend to have a higher GC content than introns (Zhu *et al.*, 2009). Indeed, one would, at first sight, expect the bias to exaggerate intronic peak density, rather than exonic peak density. However, peaks are called based on the locations of the 3' ends of reads, whereas the bias affects the locations of the 5' ends of reads. Therefore, many reads whose 3' ends are in GC-rich exonic sequence are expected to have their 5' ends in AT-rich intronic sequence.

Further examination of the meta-profiles reveals that the simulated peaks do not display the increased peak density in the beginning of the exons (first highlighted region in **Supplementary Methods Figure 2B**), as observed for true data (**Supplementary Methods Figure 2A**). It thus appears that this peak cannot be explained by read 5' end biases. However, the second highlighted region shows increased peak densities both in true and simulated data (**Supplementary Methods Figure 2A-B**). To verify if the increase observed in this region with true data could be explained by read 5' end nucleotide composition biases, we divided the true data into two classes based on whether or not any simulated peaks mapped to within the second highlighted region (**Supplementary Methods Figure 2C-D**). There was no visible difference in true mean peak densities between the two groups within the second highlighted region. This suggests that any nucleotide biases acting at the 5' ends of reads are unlikely to explain this increase in peak densities.

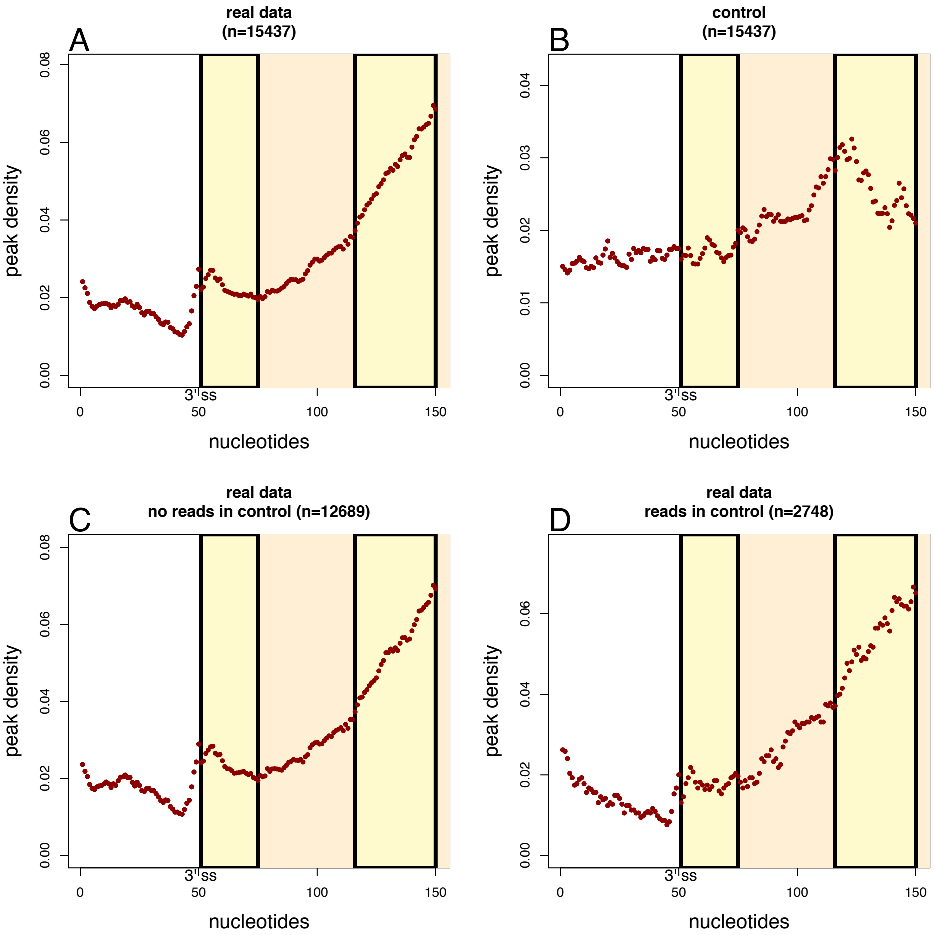


*Supplementary Methods Figure 2: average peak density for the last 50 nucleotides of introns and the first 100 nucleotides of exons (concatenated). The orange boxes show the positions of exons. Only exons at least 100 nt long have been included.*

Dingwall, C, Lomonossoff, GP, and Laskey, RA (1981). High sequence specificity of micrococcal nuclease. Nucleic Acids Res 9, 2659–2674.
Gaffney, DJ, McVicker, G, Pai, AA, Fondufe-Mittendorf, YN, Lewellen, N, Michelini, K, Widom, J, Gilad, Y, and Pritchard, JK (2012). Controls of Nucleosome Positioning in the Human Genome. PLoS Genet 8, 1–13.
Hörz, W, and Altenburger, W (1981). Sequence specific cleavage of DNA by micrococcal nuclease. Nucleic Acids Res 9, 2643–2658.
Zhu, L, Zhang, Y, Zhang, W, Yang, S, Chen, J, and Tian, D (2009). Patterns of exon-intron architecture variation of genes in eukaryotic genomes. 12, 1–12.
